# Comparative Structural and Dynamics Study of Free and gRNA-bound FnCas9 and SpCas9 Proteins

**DOI:** 10.1101/2022.03.30.486385

**Authors:** Gayatri Panda, Arjun Ray

## Abstract

The introduction of CRISPR/Cas9 based gene editing has greatly accelerated therapeutic genome editing. However, the off-target DNA cleavage by CRISPR/Cas9 protein hampers its clinical translation, thereby hindering its widespread use as a programmable genome editing tool. Although Cas9 variants with better mismatch discrimination have been developed, they have significantly lower rates of on-target DNA cleavage. Here, we have compared the dynamics of a more specific naturally occurring Cas9 from *Francisella novicida* (FnCas9) to the most widely used, SpCas9 protein. Long-scale atomistic MD simulations of free and gRNA-bound forms of both the Cas9 proteins were performed, and their domain rearrangements and binding affinity with gRNA were compared to decipher the possible reason behind the enhanced specificity of FnCas9 protein. The greater binding affinity with gRNA, high domain electrostatics, and more volatility of FnCas9 than SpCas9 may explain its increased specificity and lower tolerance for mismatches.

**Highlights:**

- The gRNA binding led to the opening of the structure of Cas9 endonuclease and exposure of hydrophobic residues to accommodate it.
- Concerted domain movement was observed in both SpCas9 (HNH-REC2 domains, CTD-Topo domains, and REC3-RuvC) and FnCas9 (HNH-REC3-REC1 and PI-WED-RuvC).
- Both Cas9s have shown a decrease in percentage helicity and an increase in the percentage of structurally dynamic residues (more secondary structural transitions) post gRNA binding.
- In the gRNA-bound form of FnCas9, the PAM-interacting domain was found to have a larger structural transition and increased coil%.
- Electrostatic interactions have a major contribution to the high binding affinity of FnCas9 with gRNA.
- Arginine-helix has shown a pivotal role in the binding of gRNA in FnCas9 protein.

## Introduction

CRISPR/Cas (clustered regularly interspaced short palindromic repeat-CRISPR associated) systems confer viral resistance by recognizing and cleaving invading nucleic acids and have been found in the genomes of most archaea, and nearly half of bacteria^1;2^. The CRISPR/Cas immune response involves three stages - spacer acquisition/adaptation, crRNA expression-processing, and interference. The CRISPR/Cas system consists of a DNA cleaving enzyme, CRISPR-associated endonuclease (Cas protein), and its associated gRNA components (preceding the Cas operon), crRNA (CRISPR RNA), and tracrRNA (trans-activating gRNA, non-coding RNA links Cas9 with crRNA) which recognize the DNA cleaving region and initiate a double-stranded break.

The most often employed DNA cleaving enzyme from the type-II CRISPR/Cas system is SpCas9, which is derived from the bacteria red *Streptococcus pyogenes.* SpCas9 enzyme is 1368aa long (4.1 kb) and adopts a bilobed architecture containing a nuclease lobe (NUC) and alpha-helical lobe or recognition (REC) lobe^3^. The nuclease lobe (residues 51-55 and 719-1368) comprise HNH (residues 780-906) and the RuvC-like domain residues (residues 1-55; 719-765; and 919-1099), Topoisomerase domain (residues 1099-1200) and C-terminal domain (residues 1201-1368). The HNH and RuvC domains help in DNA cleavage by bringing double-strand breaks into the DNA. The recognition lobe contains a ‘bridge-helix’ (residues 59-76) which connects the NUC and REC lobes with positively charged arginine residues^4;5^. Despite the rising use of the CRISPR/Cas9 technology in genome engineering, there are significant constraints that prevent it from being widely used. This includes Cas9’s large size (which makes delivery difficult), recognition of a specific PAM (which limits its efficiency), and the introduction of random off-target mutations in target gene sequences. The CRISPR/Cas system’s site-specific cleavage fails 15% of the time^6;7;8^, resulting in unintended genetic alterations and off-target effects. This limitation of Spcas9 led to the creation of novel Cas variants (e.g., Hifi Cas9, eSpCas9(1.1), SpCas9-HF1, SpCas9-NG, xCas9, Cpf1) with better efficiency^9;10;11;12^. The varying specificity and sensitivity of Cas variants emphasize the relevance of different Cas9 domains.

In work by Acharya et al., an ortholog of Cas9 from *Francisella novicida* (FnCas9), was shown to have very low non-specific editing compared to SpCas9^13^. FnCas9 consists of 1,629 amino acids and is significantly larger than other Cas9 orthologs such as SpCas9 (1,368 amino acids) and SaCas9 (1,053 amino acids). FnCas9 comprises seven domains — the REC1–3, RuvC, HNH, WED (Wedge), and PI (PAM interacting) domains. The structural comparison of FnCas9 with SpCas9 revealed unanticipated conserved and divergent features of inter-domain interaction^14^. In contrast to *Streptococcus pyogenes* Cas9 (SpCas9), which has shown variable levels of off-targeting due to tolerance of mismatches predominantly in the “non-seed” region of the gRNA, FnCas9 has been reported to have strong intrinsic specificity for its targets by tolerating only a single mismatch in the gRNA at the 5’position of the PAM distal end^14^. Acharya et al. reported the specificity of FnCas9 is determined by its minimal binding affinity with DNA in the presence of mismatches, the staggered cleavage at the cleavage site, and higher homology-directed repair rates^13^. They have also suggested the basis behind the specificity of FnCas9 could be due to the interaction of its substrate with expanded REC3 and REC2 domains which are structurally different from those in SpCas9 and SaCas9^14^. They also speculated that FnCas9’s high electrostatic potential and interactions with several bases on PAM distal and proximal ends of the substrate are necessary for FnCas9 to achieve its cleavage competent state.

Extensive structural and mechanistic studies involving atomistic simulations, targeted-MD, and accelerated-MD provided interesting insights into the understanding of the recognition mechanism of SpCas9 domains, the mode of action, and the kinetics behind SpCas9 mediated DNA cleavage. In 2016, Palermo et al. performed all-atom molecular dynamic simulations of Apo-SpCas9, SpCas9-gRNA (binary complex) state, SpCas9-gRNA-tDNA bound to incomplete ntDNA and complete ntDNA state to reveal the conformational reorganization of Cas9 associated with nucleic-acid binding. They identified that the catalytic residue H840 was at a distance of 25 Å from the scissile phosphate of the tDNA in the absence of ntDNA, whereas in the presence of ntDNA, H840 stabilizes at a smaller distance of 15 Å from the scissile phosphate of the tDNA, suggesting that ntDNA binding is crucial for the formation of a catalytically competent Cas9^15^. Gaussian Accelerated MD (~ 16 μs) on the pre-catalytic state of SpCas9 (PDB ID: 5F9R) showed how upon DNA-binding, the conformational dynamics and flexibility of the HNH domain might facilitate the unwinding of dsDNA and formation of R-loop structure triggering the formation of active state^16;17^. Ab-initio MD study of the pre-catalytic state and active-state of SpCas9 was performed to disclose the two-metal-dependent mechanism of phosphodiester bond cleavage by RuvC Active site of CRISPR/Cas9^18^.

While much has been studied for the SpCas9^15;17;18^, the cleavage and recognition mechanism of FnCas9 and the factors responsible for its higher specificity are yet to be explored. Questions pertaining to the molecular inter-play for the interactions and a comparison between the two orthologs can provide us insights into the elusive future of designing customized Cas9 molecules.

To address these questions, crystal structures of gRNA-free and gRNA-bound forms of SpCas9 and FnCas9 complexes were simulated for 1 μs and analyzed. The principal component analysis, free energy landscape, and clustering methods were used to investigate the slow motions of Cas9 proteins and conformational changes due to the binding of gRNA. The relative domain movement was studied by distance plots between the center of mass of each domain pair. The crucial interactions and key residues involved in the binding of gRNA with Cas9 protein were examined at the Cas9 protein-RNA interfaces by binding free energy calculations.

## 1 Results and Discussions

### 1.1 Conservation Analysis reveals the diversity of Sp- and FnCas9 proteins

The results from ConSurf^19^ revealed that bridge helix (BH) and the HNH domains were most conserved (SpCas9: 44% and 41%; FnCas9: 43% and 35%) and the REC2 domain was found to be least conserved (SpCas9: 12%; FnCas9: 7%) in both the proteins shown in Figure 1A. In Figure 1B, the conservation scores (Color codes) assigned by ConSurf were used to illustrate residue-wise and domain-wise conservation. Since the bridge-helix is an arginine-rich motif that bridges the nuclease and REC lobe, hence, the percentage of charged residues in this region is higher in both the proteins (SpCas9 (BH-Charged residues): 22.8% and FnCas9 (BH-Charged residues): 25%). In all the other domains, non-polar residues have shown a major involvement in conservation. Pairwise-sequence alignment of SpCas9 and FnCas9 sequences was performed using EMBOSS, which implements Needleman-Wunsch Algorithm to find the optimum alignment (including gaps) of two sequences along their entire length^20^. 23.9% residues in SpCas9 were conserved with the FnCas9 sequence (Supplementary file 2).

**Figure 1:**
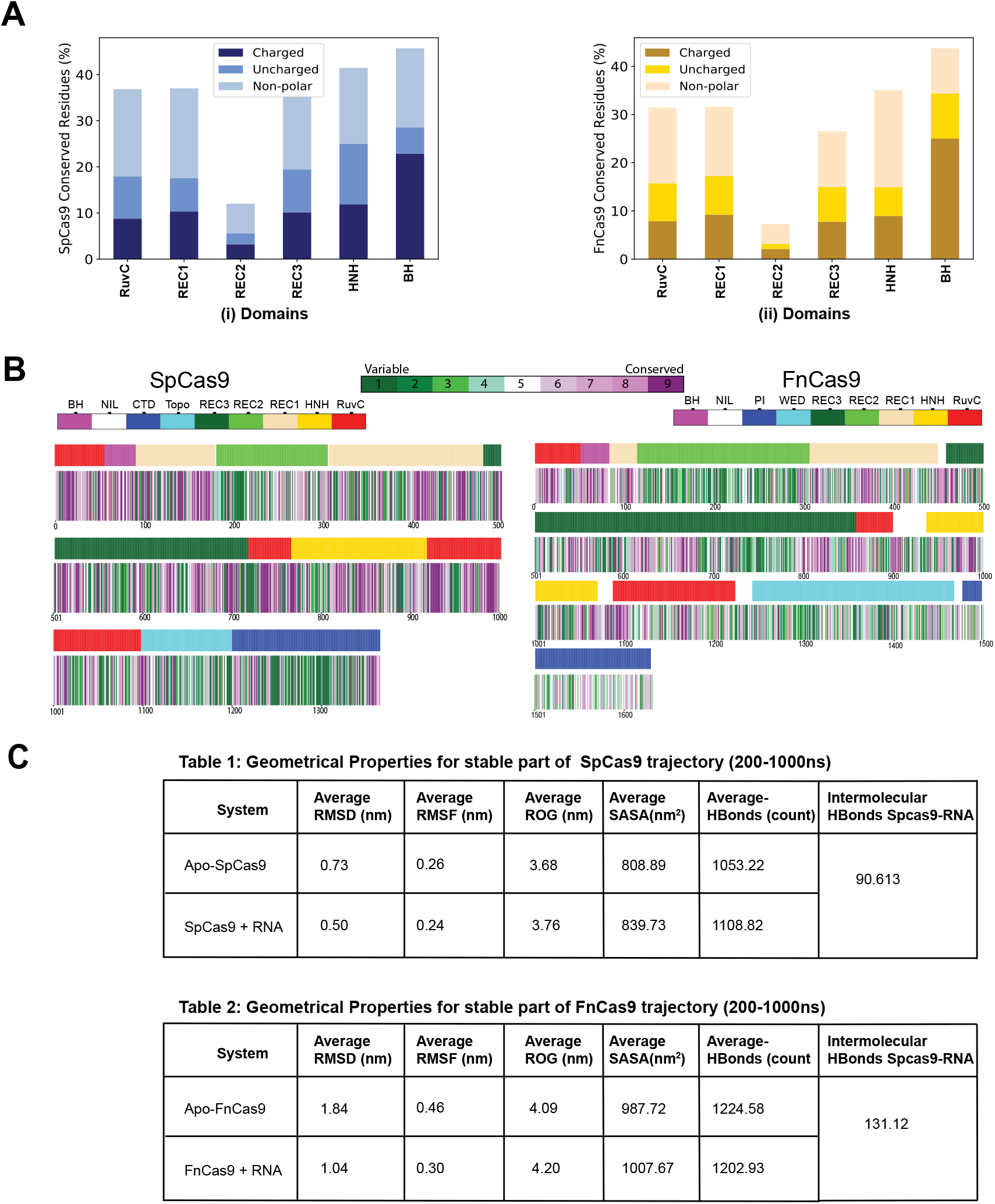
A. Grouped Bar plot showing the percentage of conserved residues (residues with color codes ≥8) in SpCas9 (PDB ID: 4CMP) (on the left) and FnCas9 (PDB ID: 5B2Q) (on the right) domains and the contribution made by charged, uncharged and non-polar residues in conservation. B. Residue and domain-wise conservation of residues in SpCas9 (on the left) and FnCas9 (on the right). The color schemes followed for domains and conservation color codes are added to the color bar on the top. C. Average geometrical properties calculated from stable part of MD trajectories for SpCas9 and FnCas9 systems.

### 1.2 gRNA-binding to Cas9 proteins stabilizes the complex

The atomic fluctuations and structural stability of the SpCas9 and FnCas9 systems during MD simulations were examined by monitoring the backbone root mean square standard deviation (RMSD) with respect to their starting structures as a function of simulation time. All the systems exhibited a stable deviation after 200 ns (200-1000 ns is considered a stable part of the trajectory). Various geometrical properties, including the number of intra-protein hydrogen bonds, solvent accessible surface area (SASA), intermolecular hydrogen bonds between Cas9 protein and gRNA, and radius of gyration, were calculated from the equilibrium MD trajectories (Supplementary Figure S1 and Supplementary Figure S2). Figure 1C list the comparison between the average values of these geometrical properties in bound and unbound form for both the Cas9 systems. Time evolution of these properties was computed (Supplementary Figure S1, Figure S2), revealing some subtle differences in geometrical properties in both cases. In the case of SpCas9, the average solvent accessibility (SASA) (Supplementary Figure S1-C) and the average number of intra-protein hydrogen (Supplementary Figure S1-D) bonds increased after gRNA-binding with a small increase in the compactness of the structure(Supplementary Figure S1-B). The SASA plot (Supplementary Figure S1-C) and average SASA values listed in Table 1 (Figure 1C) suggest an increase in solvent accessibility due to gRNA-binding leading to exposure of hydrophobic residues and loss of compactness/ folding of the protein. In the case of FnCas9, a similar observation was made with an increased radius of gyration (Supplementary Figure S2-B) and SASA after gRNA binding (Supplementary Figure S2-C), while the number of intra-protein hydrogen bonds (Supplementary Figure S2-D) decreased after gRNA-binding. The average number of intermolecular hydrogen bonds between FnCas9 protein and gRNA (94-mer) (Supplementary Figure S2-E) was more than in SpCas9 gRNA bound form (85-mer) Supplementary Figure S1-E).

**Table 1:** Computational alanine scanning result for hotspot residues involved in binding of gRNA with SpCas9 and FnCas9 protein. All values are in kcal/mol.

| HotSpot Residues | $\Delta\Delta G_{(CAS)}$ |
| --- | --- |
| Phe105 (SpCas9) | 9.95 +/- 1.2 |
| Arg55 (FnCas9) | 15.23 +/- 1.5 |
| Arg62 (FnCas9) | 13.06 +/- 0.8 |
| Arg63 (FnCas9) | 13.15 +/- 1.8 |
| Arg401 (FnCas9) | 12.87 +/- 1.7 |
| Arg455 (FnCas9) | 17.00 +/- 1.7 |

These observations suggest that gRNA-binding has stabilized the system (increased NHB); however, a reduced the compactness of the structure (increased ROG). This has led to increased solvent accessibility due to gRNA-binding and exposure to hydrophobic residues.

Residue-wise, RMSF is an often-used measure of conformational variance which highlights the regions of high mobility. To evaluate and compare the structural flexibility between free and gRNA bound forms of SpCas9 and FnCas9, per-residue root means square fluctuation (RMSF) was computed for the stable part of the trajectory (200-1000 ns) and reported in Figure 2A for SpCas9 and Figure 3A for FnCas9. In Figure 2A, we observe that the RMSF curve for Apo-SpCas9 (violet) and SpCas9 with gRNA (light-coral) had similar atomic fluctuations with average RMSF, 0.26 nm and 0.24 nm. These RMSF values were mapped onto the protein structure to find regions of higher flexibility. Regions in REC1, Topoisomerase, and RuvC-III domains were found to have higher fluctuation (RMSF >0.6 nm) in unbound form (shown in Red color in Figure 2B), whereas only the Topoisomerase domain and some portions of CTD domains were found to be highly fluctuating in gRNA bound form of SpCas9, indicating lesser fluctuating regions post-gRNA-binding.

**Figure 2:**
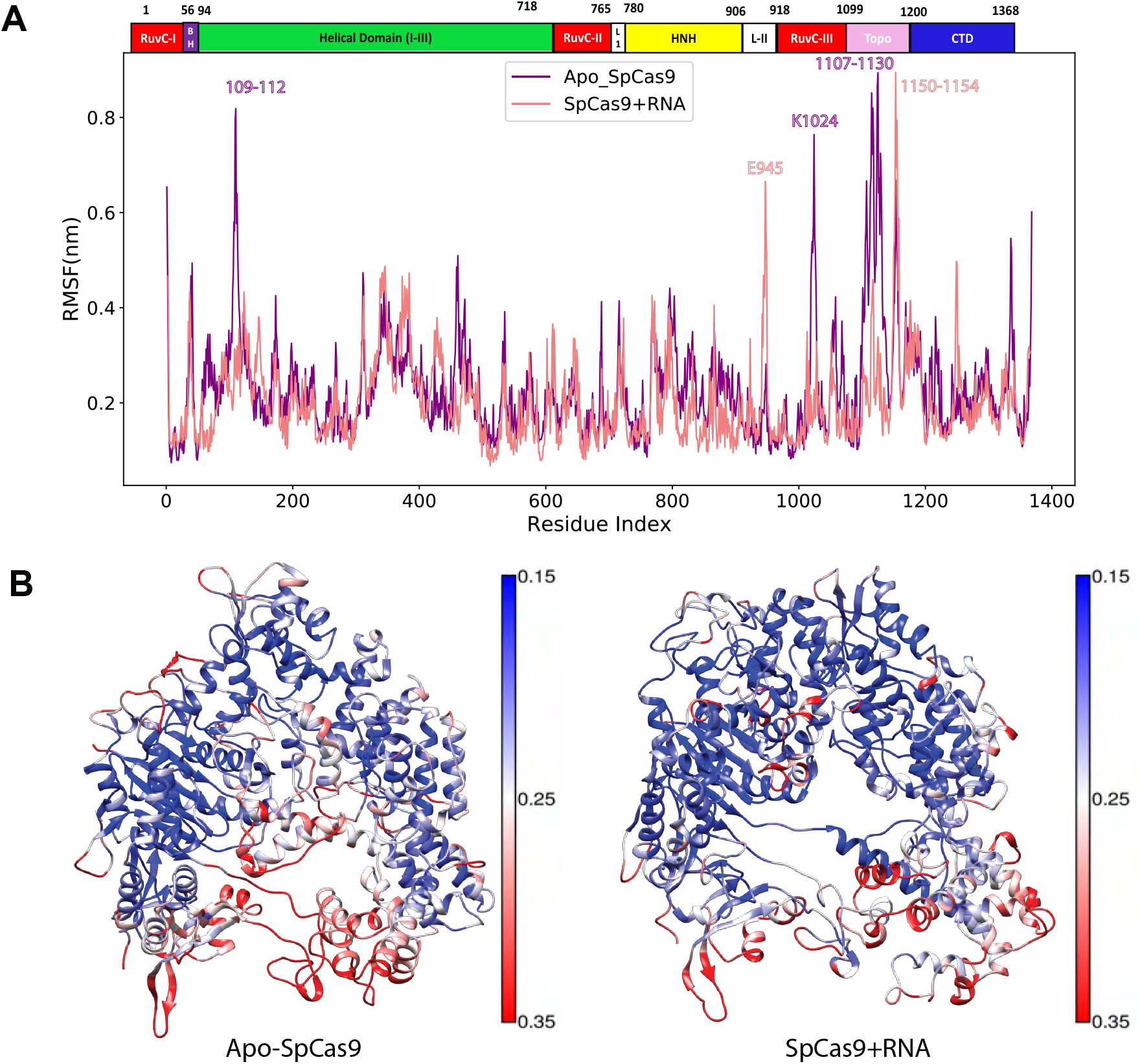
Comparison between the structural flexibility of Apo-SpCas9 and SpCas9 with gRNA. (A) Per-residue RMSF profiles were calculated from MD trajectories of Apo-SpCas9 (violet) and SpCas9+RNA (coral). The residues corresponding to every domain in the SpCas9 structure are shown. (B) The structural representations of Apo-SpCas9 and SpCas9+RNA mapped with per-residue RMSF values generated using UCSF Chimera. The color ranges from blue to red and denotes that RMSF varies from the lowest to the highest values.

**Figure 3:**
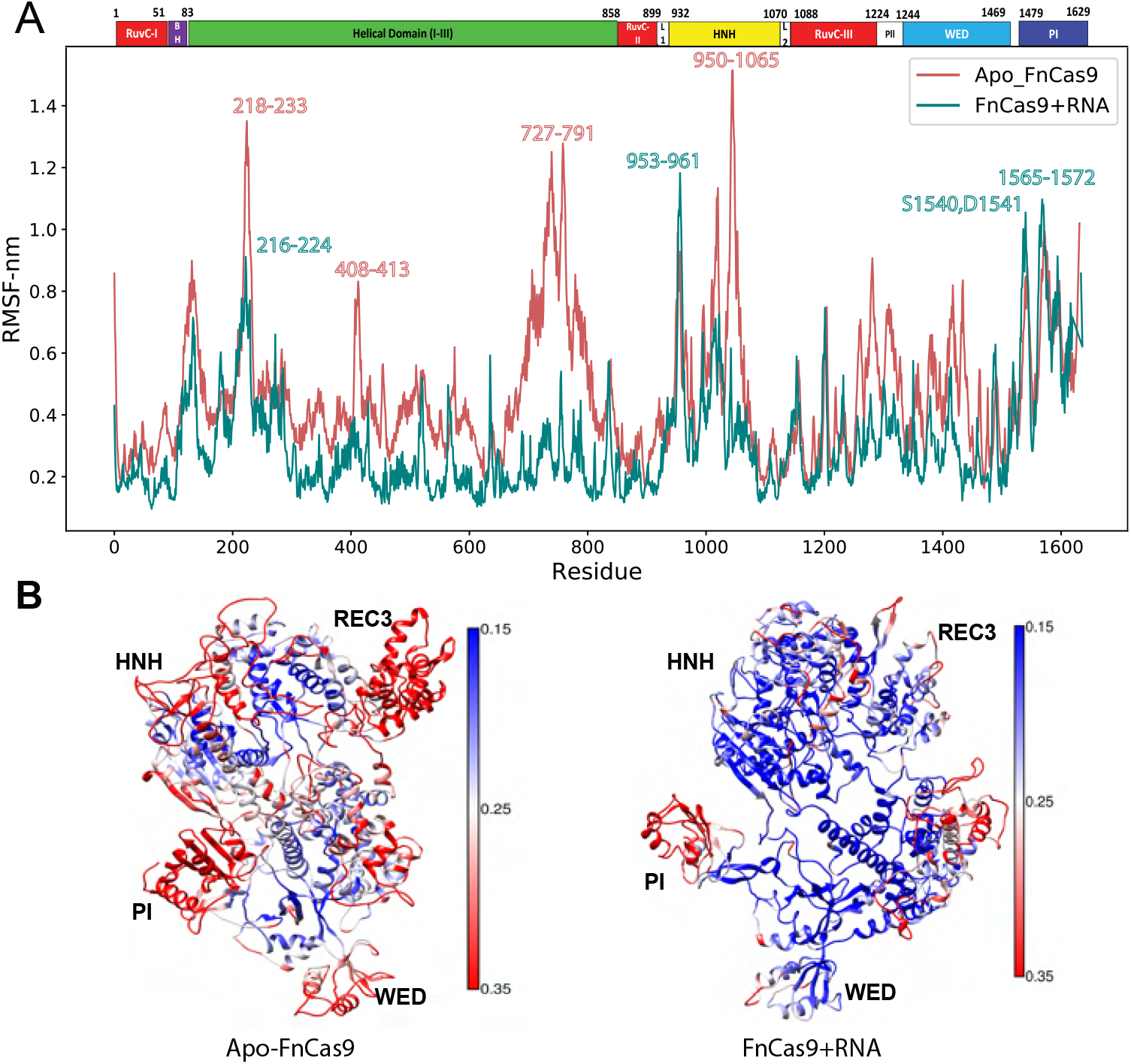
Comparison between the structural flexibility of Apo-FnCas9 and FnCas9 with gRNA. (A) Per-residue RMSF profiles were calculated from MD trajectories of Apo-FnCas9 (Indian-red) and FnCas9+RNA (teal). The residues corresponding to every domain in the FnCas9 structure are shown in the top horizontal bar. (B) The structural representations of Apo-FnCas9 and FnCas9+RNA mapped with per-residue RMSF values generated using UCSF Chimera. The color ranges from blue to red and denotes that RMSF varies from the lowest to the highest values.

The atomic fluctuations in Apo-FnCas9 (indian-red) were slightly higher than the gRNA-bound form (teal) with average RMSF, 0.46 nm and 0.30 nm (Table 2 in Figure 1C). The regions with high mobility (RMSF >0.6 nm) were determined by mapping RMSF values onto the FnCas9 protein structure. It was found that regions in the REC lobe, specifically segment of REC2-REC3 domains (residues 218-233; 727-791), HNH (residues 950-998; 1004-1065) were highly fluctuating in the free-FnCas9 form (shown in red color) and only some regions of REC2 domain (residues 216-224), HNH domain (residues 953-961; 1004-1065), and PI domains (residues 1565-1572) had high-mobility in the gRNA bound form of FnCas9 as shown in Figure 3. Both states of SpCas9 were equally flexible, whereas Apo-FnCas9 was relatively more flexible than its gRNA-bound form. On comparing, the radius of gyration, RMSD, and RMSF values, we can say that FnCas9 is more volatile than SpCas9 as it had a higher average radius of gyration and RMSF values.

### 1.3 Relative Domain Movement and Domain Structural Deformations are involved in the binding of gRNA to Cas9 proteins

Many structural and biophysical studies have reported about the important conformational changes in Cas9 protein due to nucleic acid binding^21;15;17^. Upon DNA binding, the HNH domain undergoes a significant conformational change leading to activation of Cas9 protein for catalysis of TS^22;18^. Enhanced MD simulations of pre-activated CRISPR–Cas9 (5F9R.pdb) (~ 16 μs) revealed that the tight-coupling between REC lobe and HNH domain aided in bringing the Cas9 to the activated state. The conformational transitions observed in the REC lobe disclosed how the recognition domains ‘sense’ nucleic acids, ‘regulate’ the HNH conformational change, and ultimately ‘lock’ the HNH domain at the cleavage site contributing to its catalytic activation^23^. These studies justify the significance of conformational dynamics of CRISPR–Cas9 in DNA catalysis, thereby improving its genome editing abilities.

The relative movements of intramolecular domains due to the binding of gRNA were determined by calculating distances between the center of mass of each domain as a function of simulation time for both the Cas9 forms.

The center of mass of HNH, RuvC, REC1, REC2, REC3, PI, and CTD domains were computed for both the systems (SpCas9 and FnCas9), and the distances between COM of every domain pair for the simulation length were calculated (Supplementary Figure S3 (SpCas9) and S4 (FnCas9)). The domain movement due to gRNA binding was quantified by plotting a clustermap using the difference in average domain distance (200-1000 ns) in free and gRNA-bound form. Upon gRNA binding in SpCas9, we observed that the domain movement by HNH-REC2 domains, CTD-Topo domains, and REC3-RuvC domains was similar (Figure 4A). The HNH and REC2 domains moved closer to REC3-RuvC domains and moved away from the REC1 domain after gRNA binding in SpCas9 form. Similarly, CTD and Topo moved closer to REC2 and REC3 and moved away from RuvC and REC1 domains. The RuvC and REC3 domains moved closer to REC2 and HNH domains and moved farther from Topo domains. In Fncas9, the PI, WED, and RuvC domains moved closer to each other, and REC3 and HNH domains, and moved away from the REC2 domain (Figure 4B). REC1 and REC2 domains have shown a similar domain movement by moving away from RuvC, PI, and WED domains. The FnCas9 protein has shown larger domain deviations as compared to SpCas9, as the domain deviation scale was larger in FnCas9.

**Figure 4:**
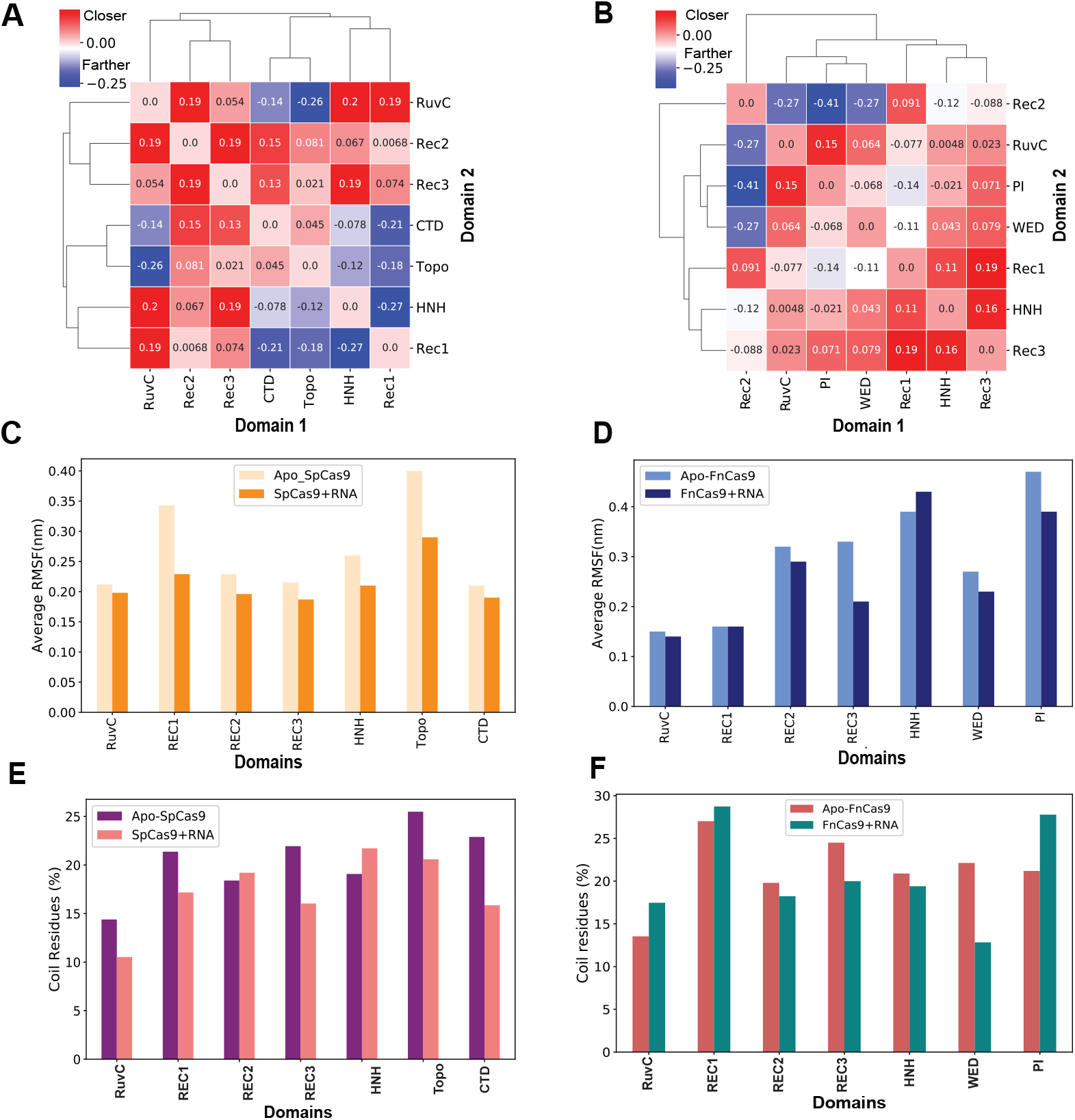
Domain Structural movements and transitions in Clustermap shows domain movement between domains on the X and Y-axis. Red boxes suggest domains that moved closer, and blue boxes are for domains that moved away. The color intensity depends on the magnitude of domain movement. (A) Clustermap for SpCas9 (B) Clustermap for FnCas9 Grouped bar plot with average RMSF values (200-1000ns) corresponding to RuvC, REC1, REC2, REC3, and HNH domains in the free and bound form of (C) SpCas9 system and (D) FnCas9 system. Domain-wise percentage of Coil residues (>70% part of trajectory) in (D) SpCas9 system and (E) FnCas9 system

The structural deformation of each of these domains in terms of domain flexibility and domain-wise structural transitions was studied by computing the domain-wise RMSF (Figure 4C and Figure 4D). The atomic fluctuations of every domain in the free form of SpCas9 were observed to be greater than in the gRNA-bound form. This observation was in agreement with other observations (in Figure 1 and Figure 2), suggesting increased stability in the presence of nucleic acids. The Topo and REC1 domains were found to be highly mobile in both the forms of SpCas9. The domain-wise average RMSF was calculated for Apo-FnCas9 and FnCas9 with gRNA for the stable part of the trajectory and plotted in Figure 4B. A similar observation was made in the case of FnCas9 too, where the atomic fluctuations of domains in the free form of FnCas9 were greater than the gRNA bound form for RuvC, REC1, and REC2 domains, whereas the average RMSF of the HNH domain was higher in gRNA bound form than in the free form. The PI and REC3 domains were highly mobile in the pre-gRNA-bound form of FnCas9.

To evaluate the conformational flexibility of these domains, the percentage of residues existing in coil conformation for more than 70% of the simulation time was calculated for every domain in Apo and gRNA-bound form for both the systems in SpCas9 form (Figure 4C) and in FnCas9 form (Figure 4D). In SpCas9, the percentage of residues existing as a coil for more than 70% times decreased after gRNA binding for all the domains except in the HNH and the REC2 domains. In FnCas9, the percentage of coils increased after gRNA-binding in RuvC, REC1, and PI domains.

The propensity of a residue to exist as a helix, sheet, turn, or coil was computed for the stable part of the trajectory. Also, dynamic residues with a high order of transition between secondary structures were annotated (See Materials and Methods for details). The secondary structure assignment of residues in Apo-SpCas9 and SpCas9 with gRNA are shown in Supplementary Figure S5-A(i) and Supplementary Figure S5-B(i), and Apo-FnCas9 and FnCas9 bound to gRNA are shown in Supplementary Figure S7-A(i) and Figure S7-B(i), respectively. The probability of residue existing as a helix, sheet, or coil for 1000 frames was calculated and plotted to see regions of structural deformations at the residue level (SpCas9-Supplementary Figure S5-A(ii) and Figure S5-B(ii); FnCas9-Supplementary Figure S7-A(ii) and Figure S7-B(ii)). The secondary structure transition of residues among several conformations of proteins contributes to the structural flexibility of the proteins. The percentage of these dynamic residues that make up each of the functional domains of both the forms of SpCas9 and FnCas9 proteins are shown in Figure 5A and Figure 5B. The percentage of dynamic residues has increased after gRNA binding for all the domains in bot Cas9s. In the case of Spcas9, residues with higher structural transitions were found to predominantly exist in the REC lobe, while the HNH domain had the least dynamic residues. The REC2 domain had shown a drastic increase in dynamic residues from 15% to 35% after gRNA binding, and on the contrary, HNH and CTD domains had shown less percentage of dynamic residues. This observation was in line with conservation analysis, where the HNH domain was the most conserved, and the REC2 domain was the least conserved. The conformational flexibility of a protein region also depends on the proportion of regions with NMR chemical shifts of a random-coil; regions that lack significantly ordered secondary structure (as determined by CD or FTIR)^24^. A similar observation was made in the case of FnCas9, the percentage of dynamic residues increased after gRNA binding. Here, the REC lobe contained the highest percentage of dynamic residues after gRNA binding.

**Figure 5:**
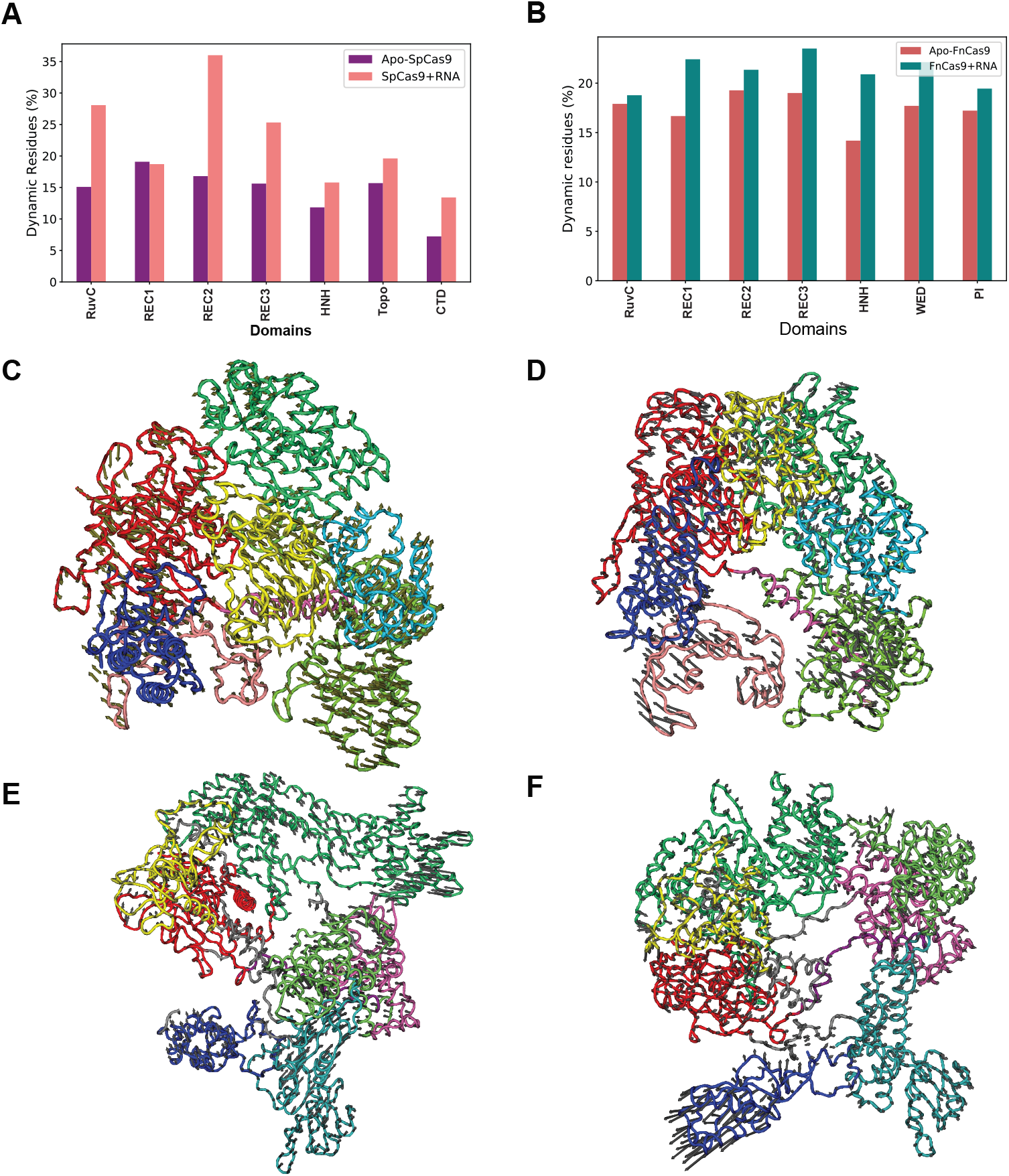
Domain Structural movements and transitions in FnCas9. Domain-wise percentage of dynamic residues (>70% part of trajectory) in (A) Apo-SpCas9 and SpCas9 with gRNA and in (B) Apo-FnCas9 and FnCas9 with gRNA. Most dominant mode, containing 100 conformations along eigenvector 1 for (C) Apo-SpCas9 and (D) SpCas9 with gRNA (E) Apo-FnCas9 (F) FnCas9 with gRNA form and visualized in NMWiz of VMD. The arrows indicate the direction and relative amplitude of motions.

After applying 70% cut-off, it was observed that the protein structure was predominantly constituted by helices (30-40% residues), followed by the dynamic residues (16-23% residues) and coils (16-20%), while sheets, turns, and bends were the least favored (4-12% respectively) (Supplementary Figure S6). Interestingly, RMSD and geometrical properties of the system during the simulation revealed that the gRNA binding was stabilizing the structure; however, the secondary structure analysis revealed a decrease in percentage helicity (Apo-SpCas9: 42.03%, SpCas9+RNA: 37.21%; Apo-FnCas9: 38.73%, FnCas9+RNA: 30.95%) and increase in the percentage of dynamic residues (Apo-SpCas9: 16.74%, SpCas9+RNA: 22.15%; Apo-FnCas9: 17.03%, FnCas9+RNA: 21.39%) due to the binding of gRNA shown in Supplementary Figure S6. The Topo and REC1 domains had higher residues as coils which could be a reason behind the high average RMSF/fluctuations (Supplementary Figure S6).

### 1.4 Larger domain deviations are observed in SpCas9 protein than in FnCas9

The principal component analysis was performed to capture the essential motions of the simulated systems. To investigate the dominant motions of free and gRNA-bound forms of Cas9 along the first principle component (PC1), corresponding to the system’s largest amplitude motion, 100 conformations of the system along eigenvector1 were stored in PDB format and visualized in Normal Mode Wizard of VMD^25^. The eigenvalues indicated fluctuations of the eigenvector in the hyperspace. Supplementary Figures S8-B (SpCas9) and Figure S8-D (FnCas9) show that only a few eigenvectors have larger eigenvalues which played a major role in the overall motion of the system.

The overlap between principal components and the coordinates of the trajectory was determined for free and bound states of SpCas9 and FnCas9. Supplementary Figure S8-A and Figure S8-C show the conformational sampling of free and bound forms of SpCas9 and FnCas9, respectively, in the essential subspace by projecting the backbone atom, showing the tertiary conformations along eigenvector 1 and 2. The 2D projection clearly depicted that free forms of Cas9 cover a wide range of phase spaces as compared to gRNA-bound forms. The decrease in the overall motion of Cas9 forms due to gRNA binding indicates the stability of the structure in the presence of nucleic acids.

In Figure 5C, the dynamics of Apo-SpCas9 protein along the first principal mode of motion (PC1) are shown. REC1, HNH, and CTD domains have shown larger domain movements. The RuvC domain moved closer to the REC lobe. The HNH and CTD domains have moved towards the REC1 domain and moved away from REC3 and REC2 domains. The dominant motions in the case of the gRNA-bound form of SpCas9 in Figure 5D included REC1, CTD, and Topo domain movements. HNH and RuvC domains moved closer to the REC1 domain. The domain movements in the case of the free form of SpCas9 were larger than the gRNA-bound form.

The most dominant mode of motion of the Apo-FnCas9 form is shown in Figure 5E. The HNH, RuvC, WED, and REC3 domains moved closer to the REC2 domain. The WED domain moved away from HNH and PI domains. In gRNA bound form (Figure 5F), major domain movement is shown by PAM interacting domain (PI domain); it moved away from the REC lobe. The reorganization of the PI domain after gRNA binding is indicative of the structural preparation of FnCas9 for the next step of gene editing, which requires scanning of the PAM motif and binding with the DNA strands. In both the Cas9 forms, the large amplitude motions were observed in the Cas9 domains, which are directly involved in the process of recognition and binding of nucleic acids, i.e., REC lobe, regions of HNH, PI, and CTD domains. The ribonucleic acid-binding has stabilized the complex. In the first mode of motion, SpCas9 protein has shown larger domain deviations as compared to FnCas9.

A three-dimensional free energy landscape (FEL) was built for the free and gRNA-bound form of both Cas9 systems using the projections of first (PC1) and second (PC2) eigenvectors to show protein stability in terms of Gibbs Free energy. Figure 6 shows the FEL values for Apo and gRNA bound form of SpCas9 and FnCas9. In the case of Apo-SpCas9, there was only one main global free energy minimum region, indicating only one stable conformational state. The lowest energy conformer of Apo-SpCas9 was identified at 636 ns with local free-energy 0 kJ/mol (State A). The second low energy state in the same energy basin had an energy of 0.74 kJ/mol (State B). The inter-conversion from State A to State B had an energy barrier of 2.2 kJ/mol, while from State B to State A had an energy barrier of 2.96 kJ/mol (Supplementary Figure S9-A). Whereas the gRNA-bound form of SpCas9 had shown two low-energy basins with energy minima at 508 ns and 850 ns (G= 0 kJ/mol). The inter-conversion between these states had an energy barrier of 3.8 kJ/mol (Supplementary Figure S9-B). The state transition barrier is slightly higher in the gRNA-bound form than in the free-form of SpCas9, suggesting that state transition is easier in Apo form than in the gRNA-bound form of SpCas9. Likewise, in Apo-FnCas9 and FnCas9 bound to gRNA, two energy basins were observed. The lowest energy conformation was extracted at 478 ns and 249 ns, respectively. In Apo-FnCas9, the lowest energy state was at 0 kJ/mol (State A), and the second low energy state had 1.6 kJ/mol (State B) of energy. The transition from State A to State B of Apo-FnCas9 had an energy barrier of 5.9 kJ/mol and from State B to State A, the energy barrier was 4.3 kJ/mol (Supplementary Figure S9-C). In the gRNA-bound form of FnCas9, the lowest energy state had 0 kJ/mol energy, and other low energy states had 0.79 kJ/mol of energy. Their inter-conversion had an energy barrier of 3.96 kJ/mol (State A to State B) and 3.17 kJ/mol (State B to State A) (Supplementary Figure S9-D). Here, energy barriers for state transitions were lower in the gRNA bound of FnCas9 form than in its free form (Supplementary S9), suggesting switch between low energy states is easier in gRNA bound form than in the Apo form of FnCas9.

**Figure 6:**
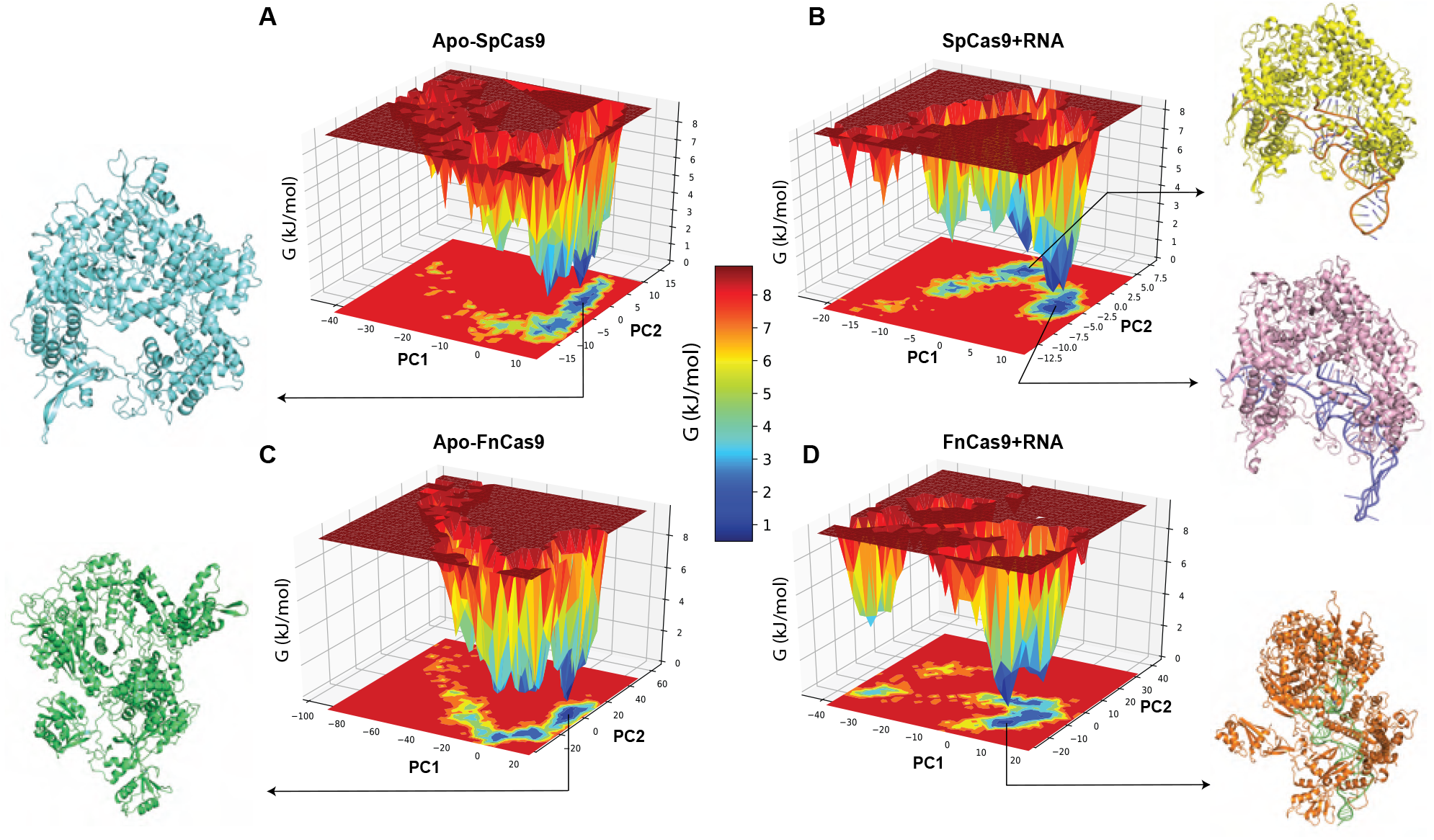
A. Free energy landscape generated using principal components (PC1 and PC2) for Apo-SpCas9 and SpCas9 with gRNA during 1000ns simulations.(B) Free energy landscape generated using principal components (PC1 and PC2) for Apo-FnCas9 and FnCas9 with gRNA during 1000ns simulations. Energy distribution is shown by the coloring pattern: Blue defines the conformational space with minimum energy (stable state) while red defines a conformational space with maximum energy (unstable state). Principal components are displayed as a contour map at the bottom of each FEL plot with similar color pattern like the energy landscape.

### 1.5 Charge potentials reveal key insights into the greater specificity of FnCas9 protein

MM-PBSA/GBSA (Molecular mechanics/Poisson–Boltzmann (Generalized-Born)) is an end-state post-processing method to estimate binding free energies. This method has been proven to balance accuracy and computational efficiency, especially when dealing with large systems. In order to calculate the binding affinity of gRNA with SpCas9 and FnCas9 forms and to understand the role of electrostatics and the atomic level interactions of gRNA to the Cas9 protein, MM-GBSA calculations were performed over an ensemble of 700 snapshots extracted from 50 ns (stable short region of the MD trajectory (950-1000 ns) using gmx _MMPBSA. Per-residue energy decomposition of MM-GBSA was used to retrieve the individual amino acid contribution to the binding energy of gRNA with Cas9 protein. The average of each energy component that contributes to the binding free energy of gRNA with SpCas9 and FnCas9 is shown in Table 7A. The guide RNA (SpCas9: 85-mer, FnCas9: 94-mer) had a lesser binding affinity with SpCas9 form (−655.4 kcal/mol) than with FnCas9 form (−1254.1 kcal/mol). In both cases, the gas phase energy, especially the electrostatic energy, had contributed predominantly to the binding energy of gRNA and protein. In the case of SpCas9 (Table 7A(i)), the contribution of electrostatic energy (−2377.7 kcal/mol) was less than the polar solvation energy (2452.0 kcal/mol). On the contrary, in FnCas9 system (Table 7A(ii)), the electrostatic interaction (−4880.2 kcal/mol) had a greater contribution than the polar solvation energy (4748.2 kcal/mol). The energetic contribution of each residue was explored to identify the hotspot residues involved in the binding of Cas9 protein with gRNA (Supplementary Figure S9). This analysis revealed that residues from the arginine-helix and REC1 domain play a pivotal role in the binding of gRNA with SpCas9 protein (Supplementary Figure S10-A(i)); however, only a few residues had shown a greater contribution to the binding energy (less than −6 kcal/mol). Many residues were found to be involved in the binding of gRNA (171) with FnCas9 protein; only those residues which have contributed more than −2 kcal/mol (< −2 kcal/mol) were plotted in Supplementary Figure S10-A(ii). These residues belonged to WED, REC3, REC1 domains, and the bridge-helix region of the protein. The contribution made by nucleotides of gRNA in binding energy was also shown in Supplementary Figure S10-B. The energy decomposition distribution for both Cas9 forms was compared in Figure 7B. The boxplot of residue-wise energy decomposition revealed that both the Cas9 forms had nearly equal median energy values with a slightly greater inter-quartile energy range for FnCas9. Interestingly, there was one outlier, Phe105 (−8.3 kcal/mol) in SpCas9 and five outliers in FnCas9, namely, Arg401 (−7.6 kcal/mol), Arg63 (−7.8 kcal/mol), Arg62 (−8.6 kcal/mol), Arg55 (−10.7 kcal/mol), and Arg455 (−12.8 kcal/mol) with energy contribution greater than −7.5 kcal/mol. These residues were highlighted on the surface of SpCas9 and FnCas9 structures in sticks representation in Figure 7D. In FnCas9, all the outlier residues (energy < −7.5 kcal/mol) were arginines, three of which were a part of the bridge helix suggesting the pivotal role of arginine helix in gRNA recognition and binding.

**Figure 7:**
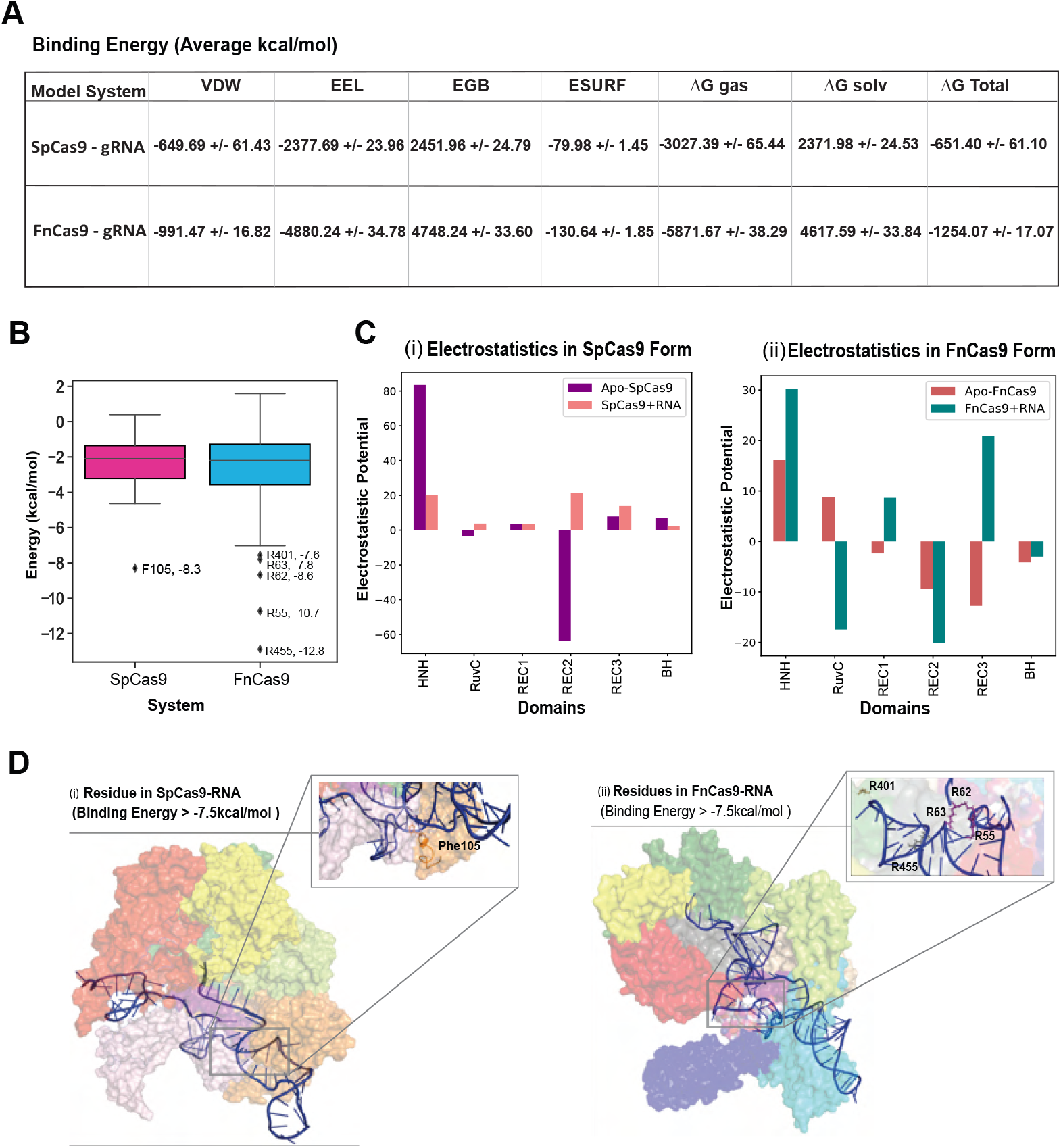
A. Bar plots indicating the free energy components (vdW, electrostatic, polar surface, SASA, obtained from MM-PBSA calculations for gRNA binding with SpCas9 (left) and FnCas9 (right). B. Box plot showing the distribution of per-residue energy contribution in the binding of gRNA with SpCas9 and FnCas9. C. Electrostatic potential of HNH, RuvC, REC domains, and bridge-helix in SpCas9 and FnCas9. D. Hotspot residues involved in gRNA binding shown in sticks representation in SpCas9 and FnCas9.

Due to the strong negative electrostatic field associated with the gRNA phosphate backbone, charged interactions are expected to be critical for correct protein-RNA binding and functioning. The MMGBSA results emphasized the significance of positively charged arginine residues located in close proximity to the gRNA-binding site. This observation can be correlated with the role of electrostatic interactions in gRNA binding. To elucidate the contribution made by each Cas9 domain in maintaining a suitable environment for gRNA binding, the electrostatic potential of each domain was evaluated using the APBS^26^ tool and compared with their gRNA-free and gRNA-bound forms in Figure 6C. A highly-positive electrostatic potential was shown by the HNH domain, and a highly-negative electrostatic potential was shown by the REC2 domain in the case of Apo-SpCas9. A drastic change in electrostatic potential was observed in the case of the REC2 domain of SpCas9 after gRNA binding, where the negative potential transformed to a positive potential. In the case of FnCas9, the RuvC domain transformed its potential from electropositive to electronegative. On comparing the electrostatics of domain centers after gRNA binding, it was found that FnCas9 domains had high electrostatic potential as compared to SpCas9 (Positive potential in HNH, REC1, and REC3 domain and Negative potential in RuvC and REC2 domains), supporting the observation made in work by Acharya et al.^13^.

Computational Alanine Scanning is an effective method to determine the significance of side-chain functional groups of hotspot residues in protein-gRNA binding by substituting it to Alanine and then evaluating the change in binding affinity between Cas9 protein and gRNA in both the systems. The binding free-energy change due to Alanine mutation of hotspot residues (Spcas9 System: Phe105; FnCas9 System: Arg401, Arg63, Arg62, Arg55, and Arg455) was performed to assess their significance in gRNA binding. The ΔΔG _(CAS)_ for these residues is shown in Table 3. The alanine conversion of Phe105 decreased the binding affinity of gRNA with SpCas9 by 9.958 kcal/mol, and Arg455 decreased the binding affinity of gRNA with FnCas9 by 17 kcal/mol (similar values were observed for other hotspot residues), indicating an unfavorable alanine-substitution demonstrating the significance of these residues in gRNA binding.

## Conclusions

Comparative analyses of gRNA-free and gRNA-bound forms of SpCas9 and FnCas9 suggested that the binding of gRNA has brought structural stability to the system. gRNA-binding has led to a decrease in structural compactness, indicating the opening of the structure and exposure of hydrophobic residues (1). The RMSF profile for both states of SpCas9 had similar flexibility. Interestingly, in FnCas9, the Apo-form was more flexible than the gRNA-bound form (2). Residues in the REC2-REC3 domains (residues 218-233; 727-791) and the HNH (residues 950-998; 1004-1065) domain were highly fluctuating in the gRNA-free form, whereas the PI domain (residues 1565-1572) had high-mobility in the gRNA bound form of FnCas9. The FnCas9 was more volatile than the SpCas9 form, shown by the high radius of gyration, RMSD, and RMSF values. Domains with higher fluctuations had shown larger domain movement, suggesting a correlation between them (especially in Apo-FnCas9). As a consequence of gRNA-binding, a concerted series of domain movements occurred in SpCas9 (HNH-REC2, CTD-Topo, and REC3-RuvC) and FnCas9 (HNH-REC3-REC1 and PI-WED-RuvC). The FnCas9 form had shown larger domain deviations as compared to SpCas9 as their difference in average domain distance range was higher than the latter. The increase in the percentage of dynamic residues observed in both Cas9s post-gRNA-binding implies the occurrence of higher structural deformations in those regions (Supplementary Figure 5 and Supplementary Figure 7). In SpCas9, the REC2 domain contained a higher percentage of dynamic residues, and HNH had a low dynamic residue percentage. This observation was in line with conservation analysis, where the HNH domain was most conserved, and the REC2 domain was least conserved. The high fluctuations and movement shown by the PI domain in the gRNA-bound form of FnCas9 could be due to higher structural transitions and increased coil percentage after gRNA-binding. In the free-energy landscape, clusters with frames of the initial part of the trajectory were structurally and energetically distinct. Energetically more stable states were observed in the gRNA-bound form in both the Cas9s. The binding free energy analysis revealed that gRNA is more tightly bound in FnCas9 form than in SpCas9. In SpCas9, the polar-solvation energy (EGB) had contributed predominantly to the binding energy, whereas in FnCas9 electrostatic energy (EEL) was the major contributor, disclosing the significance of electrostatic interactions in gRNA-FnCas9 binding. The per-residue energy decomposition in SpCas9 form revealed the significance of arginine-helix and REC1 domain in gRNA binding. Many residues have shown contribution towards the binding of gRNA in FnCas9, mainly belonging to WED, REC3, REC1 domains, and the bridge-helix region (Supplementary Figure S9). Hotspot-residues (energy < −7.5 kcal/mol) with the greatest contribution towards gRNA binding were Phe105 in SpCas9 and Arg455, Arg401, Arg55, Arg62, and Arg63 in FnCas9. All hotspot residues were arginines in the case of FnCas9, three of which were a part of the bridge helix (R55, R62, R63), suggesting its pivotal role in gRNA recognition and binding.

This study talks about the sequential differences as well as changes in dynamics at the atomic level between Cas9 orthologs, SpCas9, and FnCas9. It sheds light on the role of bridge-helix residues and electrostatic interactions in FnCas9-gRNA binding. Cas9 domains play an essential role in bringing the protein into the catalytic-competent stage and, ultimately, in the DNA cleavage process. The structural changes and differences in domain rearrangement in both Cas9 orthologs show that the cleavage mechanism differs in these systems, which might explain why Cas9s have different specificity and sensitivity. To gain a better understanding of the cleavage mechanism at the atomic level DNA bound ternary complex of SpCas9 and FnCas9 should be simulated and compared. Future research should focus on comparing the domain rearrangement and change in binding free energies in the presence of DNA and implementing high-level quantum-mechanical simulations to describe the formation and breaking of bonds to understand the DNA cleavage mechanism of FnCas9 systems, which will aid in optimizing Cas9’s activity.

## Materials and Methods

### Estimation of evolutionary conservation of Cas9 domains

The amino-acid positions which evolve slowly are commonly conserved sites that are important for protein structure and function. Proteins alter their secondary structure due to a change of surroundings or as a result of interaction with other proteins, but these conserved residues preserve their structures in all the conformations of the protein and therefore contribute less to the structural flexibility of the protein^27^. The ConSurf server^19^ was used to estimate the level of conservation of amino acid positions in different domains of the SpCas9 and FnCas9 protein based on phylogenetic relationships between their homologous sequences. The protein structure of SpCas9 (PDB ID: 4CMQ) and FnCas9 (PDB ID: 5B2Q) were submitted to the ConSurf server, which calculated the conservation scores partitioned into the discrete scale of nine bins. The position with bins >8 scores is considered conserved sites, while the amino-acid positions with bin *leq* 2 are considered variable sites. Applying these cut-offs for indicating conserved and variable residues, the percentages of conserved and variable residues were calculated for HNH, RuvC, BH, REC1, REC2, and REC3 domains in both the proteins. The proportion of charged (H, K, R, D, E), uncharged (C, S, T, Y, Q), and non-polar (A, G, I, L, M, W, F, P, V) residues in conservation was also calculated.

### System setup and MD Simulations

MD simulations have been performed using four model systems of Cas9, which were the Apo form of SpCas9 and in complex with gRNA, and the Apo form of FnCas9 and in complex with gRNA. These model systems have been prepared using the crystallographic coordinates of *Streptococcus pyogenes* Apo-SpCas9 (PDB ID: 4CMQ)^28^, SpCas9+RNA (PDB ID: 4ZT0)^16^, *Francsella novicida* Apo-FnCas9 (PDB ID: 5B2Q w/o nucleic acids)^14^ and FnCas9+RNA (5B2Q, DNA strand removed), solved at 3.09 Å, 2.90 A, and 1.70 A resolution, respectively. The four independent MD simulations were performed using GROMACS 2020 software package^29^. AMBERff99SB and AMBER94 forcefields were used for protein, and nucleic acid parameterization^30^. The system was neutralized and solvated by explicit water molecules, which were modeled by the TIP3P) parameter set^31^ in a dodecahedron box. Particle mesh Ewald was used to treat long-range electrostatic interactions^32^. The pressure and temperature were controlled with the Parrinello-Rahman coupling algorithm, and Langevin thermostat^33^. The energy minimization of the whole system was done using the steepest descent algorithm until the maximum force was less than 1000 kJ mol^-^1nm^-^1 to remove any steric clashes. The system was heated to 300 K and equilibrated for 100 ps in both Isothermal-Isobaric and canonical ensembles. The positions of Cas9 protein and gRNA were restrained with a harmonic constant of 0.1 kcal/mol A^-^2 to avoid improper geometry. A non-restraint MD simulation was performed for 1000 ns at constant 1 atm pressure. The cut-off value for nonbonded interactions was set to 12 Å. The MD trajectory snapshot was saved after every 1 ns; thus, 1000 frames for every simulation were collected for further analysis.

### Distance between Domains

The center of mass of HNH, RuvC, REC1, REC2, and REC3 domains in both the Cas9 forms was calculated using the pdb_centermass script of pdbtools (https://github.com/harmslab/pdbtools). Additionally, the COM of Topo and CTD in SpCas9 and PI and WED domains in FnCas9 protein were also determined using the same script. To quantify and compare the domain movement due to gRNA binding, the distance between the center of mass of every domain pair during the simulation time (200-1000ns, stable part of trajectory) was calculated to characterize relative movements of intramolecular domains (due to gRNA binding) as a function of simulation time. Interdomain distances in both free and gRNA-bound forms for both the systems were averaged. The difference (Dist-Free form - Dist-bound form) in these averaged distances was determined for every domain pair. A positive delta indicates that the domains came closer and vice-versa. Normalized delta distances were taken as input to form a clustermap to illustrate the relative movement of domains due to the binding of gRNA. A clustermap uses hierarchical agglomeration clustering to cluster similar columns together. The order of clustering forms a dendrogram. These values were standard scaled to 1 to make a comparison between both Cas9s.

### Domain structural transitions and fluctuations

The propensity of a residue to attain a particular secondary structural element was computed for 1000 frames (Fig S4). A secondary structure was assigned to those residues which had a greater than 70% probability of existing either as a helix, turn, strand, or coil throughout the simulation. Those residues that did not conform to any particular secondary structure throughout our simulation (existed as a secondary structure for < 70% of times throughout simulation) were also quantified and referred to as dynamic residues. The percentage of these dynamical residues that make up each of the functional domains of the protein. To determine regions with high flexibility, the percentage of residues existing in coil conformation in more than 70% of the simulation time was calculated for every domain in Apo and gRNA-bound form for both the systems. Domain-wise atomic fluctuations were studied by plotting root-mean-square fluctuation for every domain. The average RMSF of SpCas9 and FnCas9 proteins in free and bound forms were plotted.

### Principal component analysis

Principal component analysis (PCA) is a useful method for assessing trajectory data from MD simulations and identifying the essential dynamics^34;35^. Based on the calculation of the covariance matrix, this method applies a linear transformation to project the original high-dimensional representation of macromolecular dynamics into the low-dimensional space. The covariance matrix of the protein backbone atoms was built and diagonalized using gmx covar method of GROMACS 5.1 package^29^ to investigate the dominant motions of free and gRNA bound forms of Cas9. Only a very small number of eigenvectors (modes of fluctuation) contribute significantly to the overall motion of the protein. This can be seen from the eigenvalues contained in eigenval.xvg. The first principle component (PC1) corresponds to the system’s largest amplitude motion, and the system dynamics along the PC1 are generally called “Essential Dynamics”. The most dominant mode, containing 100 conformations along eigenvector 1, was saved in PDB format and visualized in Normal Mode Wizard of VMD^25^.

### MMGBSA Calculations

The Molecular Mechanics/Generalized Born Surface Area (MM/GBSA) method^36^ is a popular technique for determining binding free energies between proteins and ligands. gmx_MMPBSA^37^ was used to calculate the enthalpy contribution to binding free energy. The GB (OBC) model (igb=2) was used to calculate the solvation energy. A total of 700 snapshots, equally distributed in the last 50ns of the SpCas9+RNA and FnCas9+RNA trajectory, were selected to calculate the enthalpy contribution using MM/GBSA. The single trajectory approach was used to calculate the binding free energy. Molecular mechanics potential energy (electrostatic and van der Waals energies), the free energy of solvation (polar and nonpolar solvation energies), and the energetic contribution of each residue and nucleotide to total binding energy were obtained for both SpCas9 and FnCas9 proteins with gRNA. The gas-phase energies calculated using the Molecular Mechanics force field are a sum of van der Waals (VDWAALS) and electrostatic (EEL) energies, whereas the solvation energy includes a polar and a non-polar component. The polar component is calculated using the implicit Solvent Model, i.e. Generalized Born (GB), (EGB) and the non-Polar component depends on solvent accessible surface area (ESURF). Free energies of solvation are estimated by applying Poisson-Boltzmann (PB) calculations for the electrostatic contribution and a surface-area-dependent term for the non-electrostatic contribution to solvation.

### Electrostatic Calculation of Cas9 domains

The electrostatic contribution made by HNH, RuvC, REC1, REC2, and REC3 domains were calculated using the Adaptive Poisson-Boltzmann Solver (APBS)^38^. Prior to using APBS, the standalone version of PDB2PQR^39^ was used to assign charge and radius parameters for the CHARMM force field to all the domain structures. APBS uses iterative solvers to solve the nonlinear algebraic equations resulting from the discretized Poisson-Boltzmann equation with a fixed error tolerance of 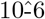. The electrostatic values for the centroid coordinates of each of these domains were then extracted from its corresponding electrostatic potential file using the multivalue script (provided in APBS).

### Gibbs Free energy landscape

The free energy landscape (FEL) is used for describing conformation exchange in biomolecular processes such as molecular recognition, folding, and aggregation. The population and stability of functionally different states are determined by free energy basins and their depths, while the inter-basin barriers correspond to the transient states that connect them. gmx sham module of GROMACS was used to compute Gibbs Free Energy using the Boltzmann equation. The relative free energy between two states is given by;

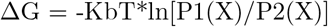

where Kb is the Boltzmann constant, T is absolute temperature, X stands for the reaction coordinate and P(X) is the probability distribution of the system along the reaction coordinate. Based on the PCA calculation, PC1 and PC2 were chosen as reaction coordinates in this analysis. Frame-wise free energy profiles of all four Cas9s are shown in Supplementary Figure S9.

### Computational Alanine Scanning

Computational Alanine Scanning is an energy-based method to calculate the effects of alanine mutations on the binding free energy of a protein-gRNA complex. This alanine substitution removes the side chain from the selected residue, which aids in the assessment of the role of a side chain functional group at a particular position. It is used to determine the significance of a residue in protein-gRNA binding. The principle followed is:

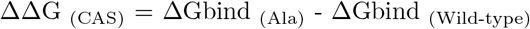

In this formula, ΔΔG_(CAS)_ is the binding free energy difference between ΔGbind_(Ala)_ (residue substituted with Alanine) and ΔGbind _(Wild-type)_. This analysis was performed to evaluate the potential of hotspot residues in gRNA binding. 50 frames from the last 10 ns part of the trajectory were considered as input for both systems.

## Acknowledgements

We would like to thank Dr. Arul Murugan (Indraprastha Institute of Information Technology, Delhi) for his critical insights and valuable suggestions in manuscript writing. We would also like to thank Dr. Debajyoti Chakraborty (CSIR-Institute of Genomics and Integrative Biology) for reviewing the manuscript. The authors would like to thank the computational facility at IIIT Delhi for supporting this work.

## Author Contribution

G.P. was involved in the acquisition of data, analysis and/or interpretation of data, drafting the manuscript, and revising the manuscript critically for important intellectual content. A.R. led the conception and design of the study, analysis, and interpretation of the data, preparing the final manuscript and its revision.

## Notes

### Competing Interest Statement

The authors have declared no competing interest.

### Summary of Updates

This is the revised version of the manuscript.

