## Supplementary File 1 for "Comparative Structural and Dynamics Study of Free and gRNA-bound FnCas9 and SpCas9 Proteins"

### Comparative Structural Dynamic Study of Free and RNA-bound state of FnCas9 and SpCas9 Proteins

### Supplementary Information

##### Supplementary Figures

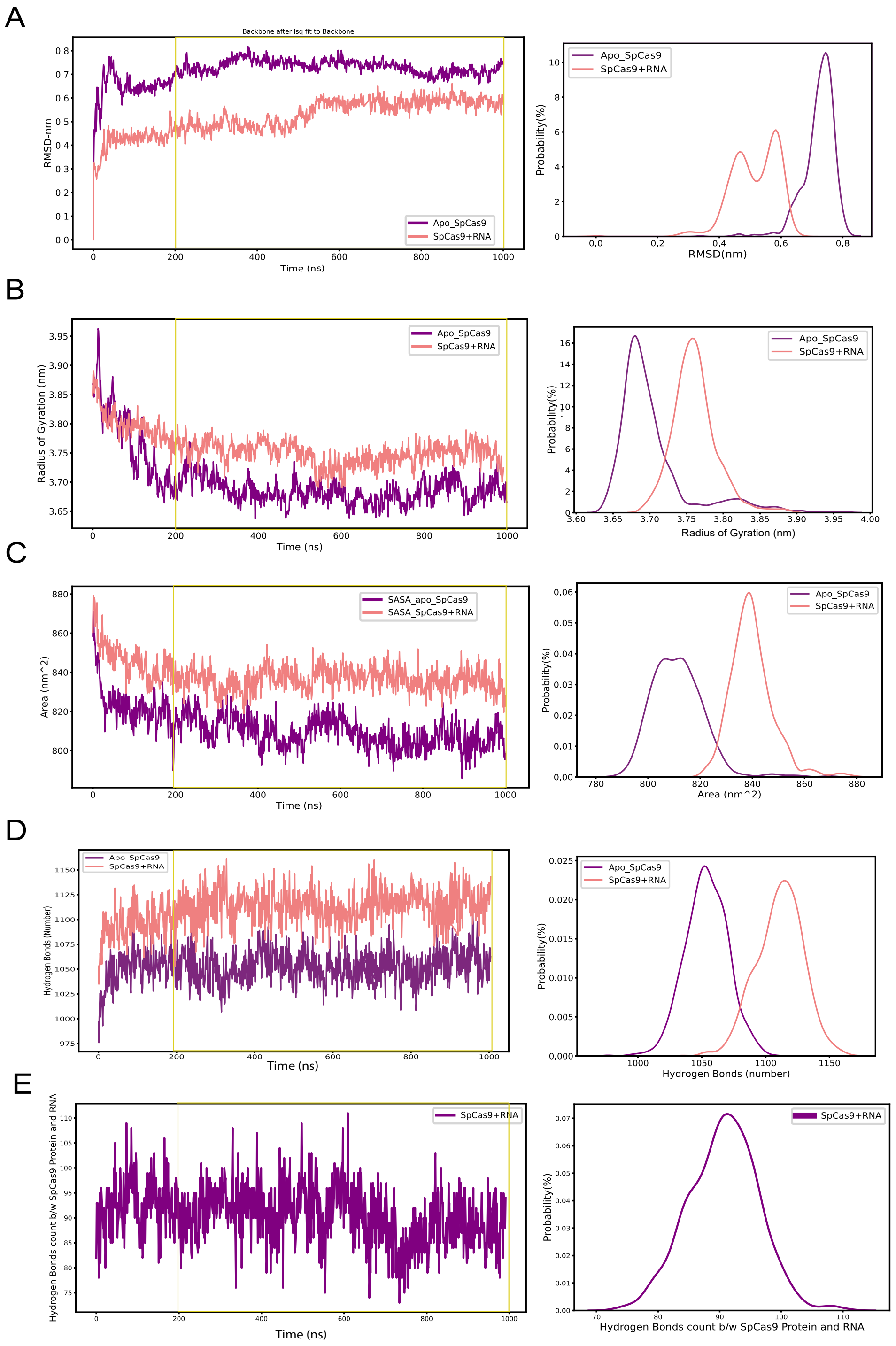


Figure S1: A.[Left] RMSDs of backbone atoms versus simulation time for Apo-SpCas9 (violet) and SpCas9 bound to RNA (light-coral). [Right] The probability distribution of RMSDs of Apo-SpCas9 (violet) and SpCas9 bound to RNA (light-coral) B. [Left] The time evolution of Radius of Gyration of SpCas9 protein (free and RNA bound form). [Right] The probability distribution of radius of gyrations of Apo-SpCas9 (violet) and SpCas9 bound to RNA (light-coral). C [Left] The solvent accessible surface area for RNA-free (violet). [Right] The probability distribution of SASA for gRNA-free and gRNA-bound forms of SpCas9 (light-coral). D. [Left] The time evolution of the number of intra-protein hydrogen bonds for Apo-SpCas9 (violet) and Spcas9 with RNA (light-coral). [Right] The probability distribution of intra-protein hydrogen bonds count for RNA-free and RNA-bound form of SpCas9 (light-coral)

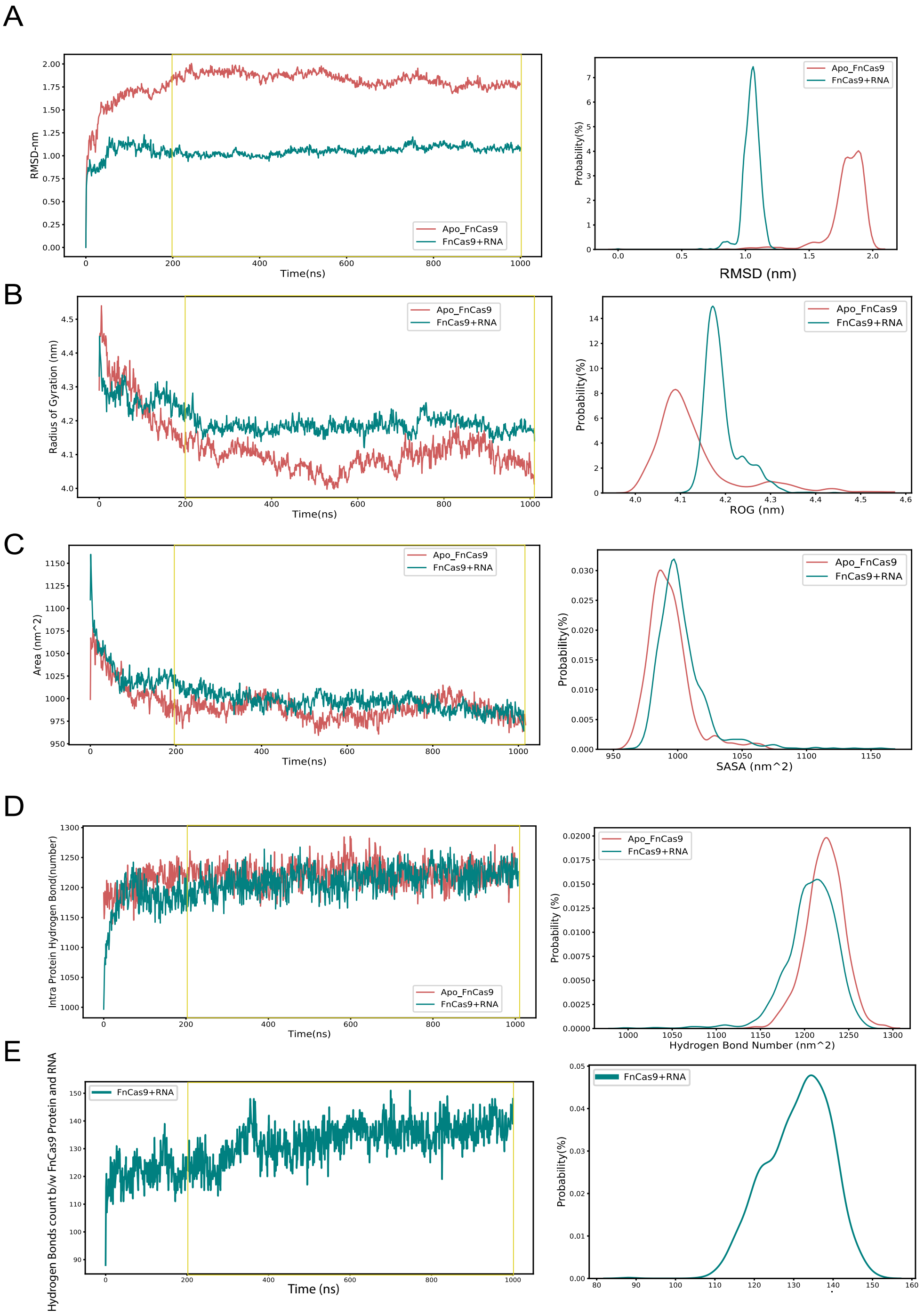


Figure S2: A.[Left] RMSDs of backbone atoms versus simulation time for Apo-SpCas9(violet) and SpCas9 bound to gRNA (light-coral). [Right] The probability distribution of RMSDs of Apo-SpCas9 (violet) and SpCas9 bound to gRNA (light-coral) B. [Left] The time evolution of Radius of Gyration of SpCas9 protein (free and gRNA bound form). [Right] The probability distribution of radius of gyrations of Apo-SpCas9 (violet) and SpCas9 bound to gRNA (light-coral). C [Left] The solvent accessible surface area for gRNA-free (violet). [Right] The probability distribution of SASA for gRNA-free and gRNA-bound form of SpCas9 (light-coral). D [Left] The time evolution of number of intra-protein hydrogen bonds for Apo-SpCas9 (violet) and Spcas9 with gRNA (light-coral). [Right] The probability distribution of intra-protein hydrogen bonds count for gRNA-free and gRNA-bound form of SpCas9 (light-coral)

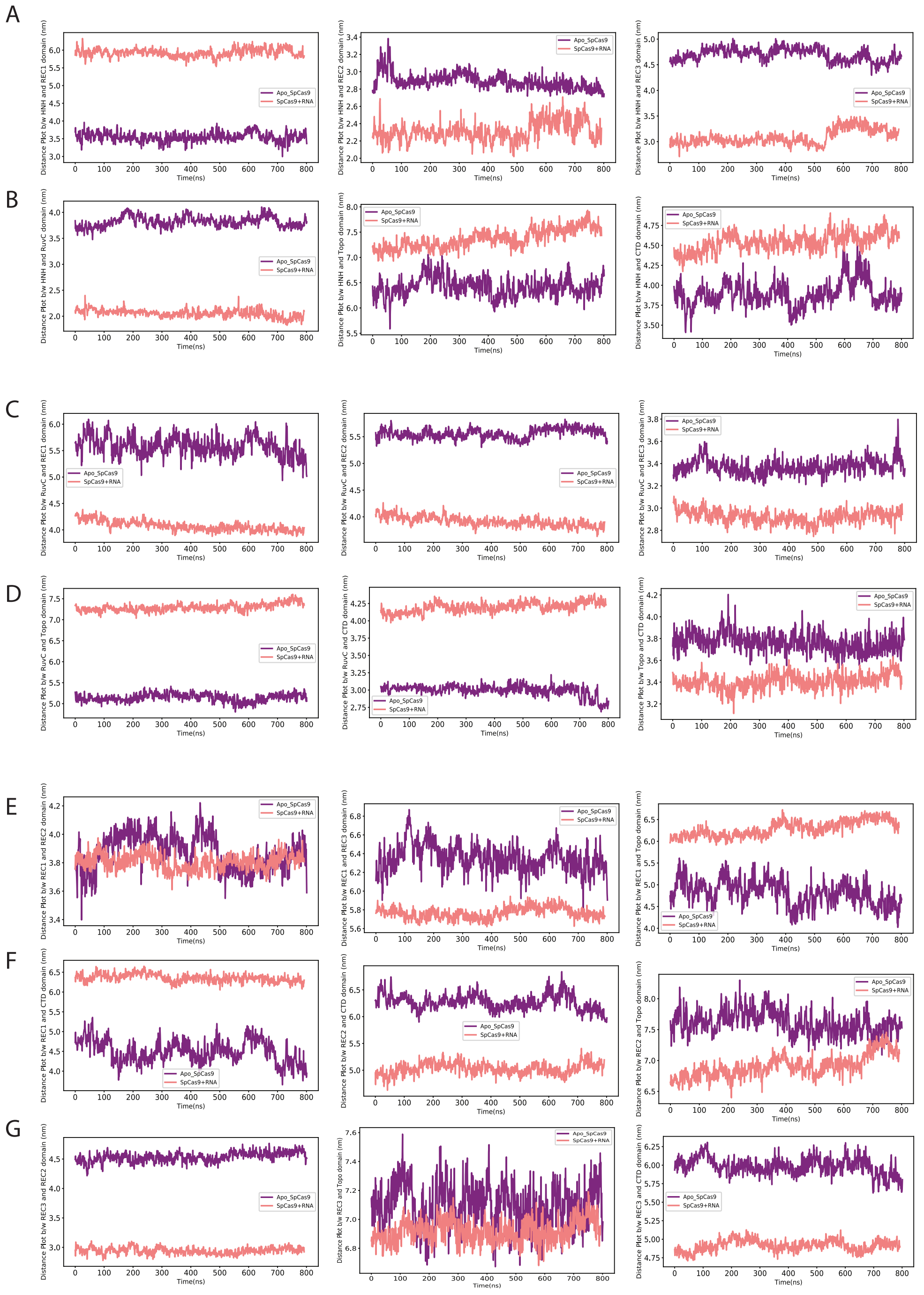


Figure S3: Distance plots for domain pairs in Apo-SpCas9 and SpCas9 with gRNA for equilibrium trajectory (200-1000 ns, duration 800 ns). Row A and Row B contain distance plots of HNH domain with RuvC, REC1, REC2, REC3, Topo and CTD domains. Row C and Row D contain distance plots of RuvC domain with REC1, REC2, REC3, Topo, CTD domains and distance plot of Topo with CTD domain. Row E contains distance plots for REC1-REC2, REC1-REC3, REC1-Topo, REC1-CTD, REC2-CTD and REC2-Topo. Row F contains distance plots for REC3-REC2, REC3-Topo and REC3-CTD domain pair.

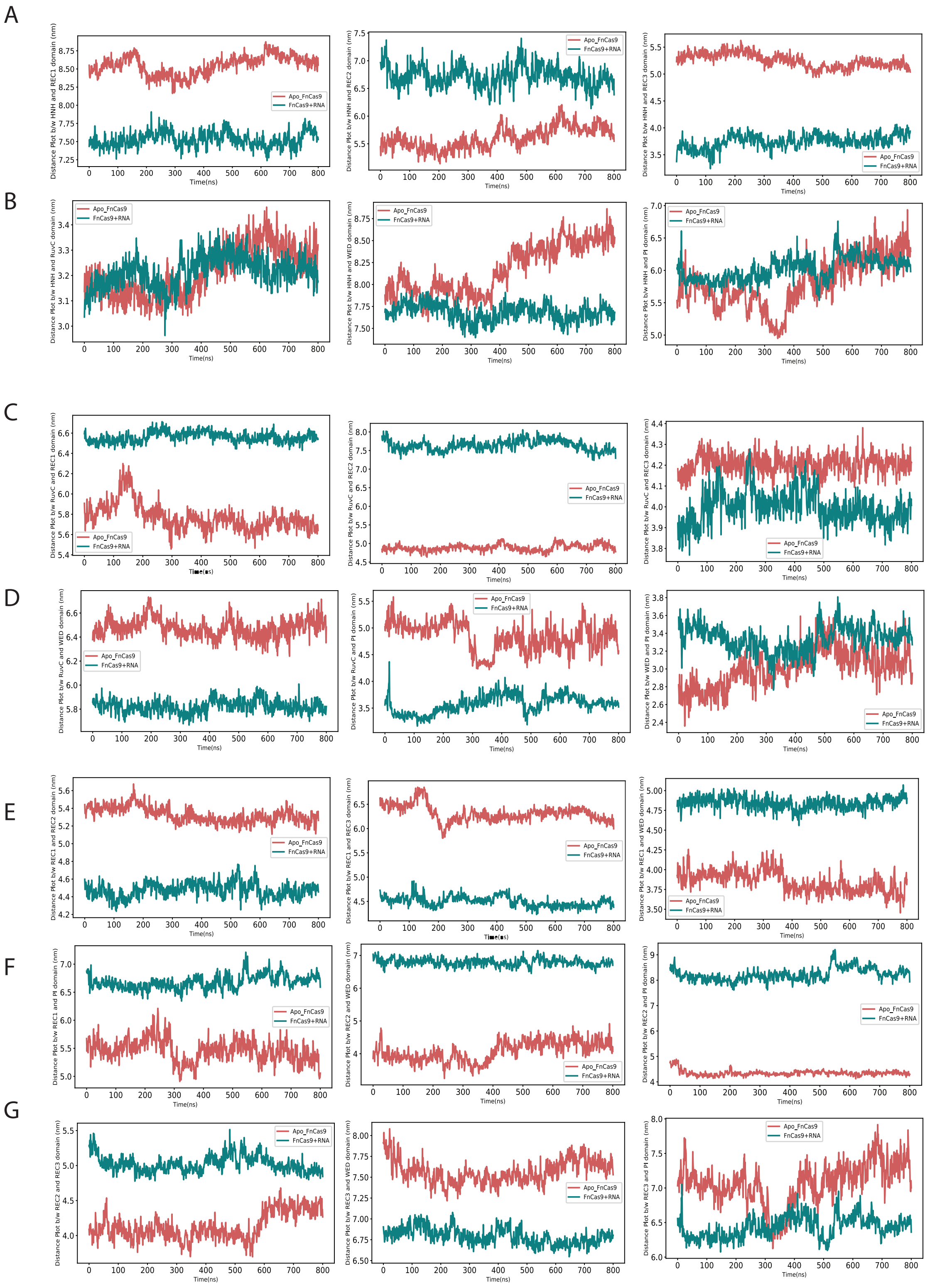


Figure S4: Distance plots for domain pairs in Apo-FnCas9 and FnCas9 with RNA for equilibrium trajectory (200-1000 ns, duration 800 ns). Row A and Row B contain distance plots of HNH domain with RuvC, REC1, REC2, REC3, WED and PI domains. Row C and Row D contain distance plots of RuvC domain with REC1, REC2, REC3, WED, PI domains and distance plot of PI with WED domain. Row E contains distance plots for REC1-REC2, REC1-REC3, REC1-WED, REC1-PI, REC2-PI and REC2-WED. Row F contains distance plots for REC3-REC2, REC3-WED and REC3-PI domain pairs.

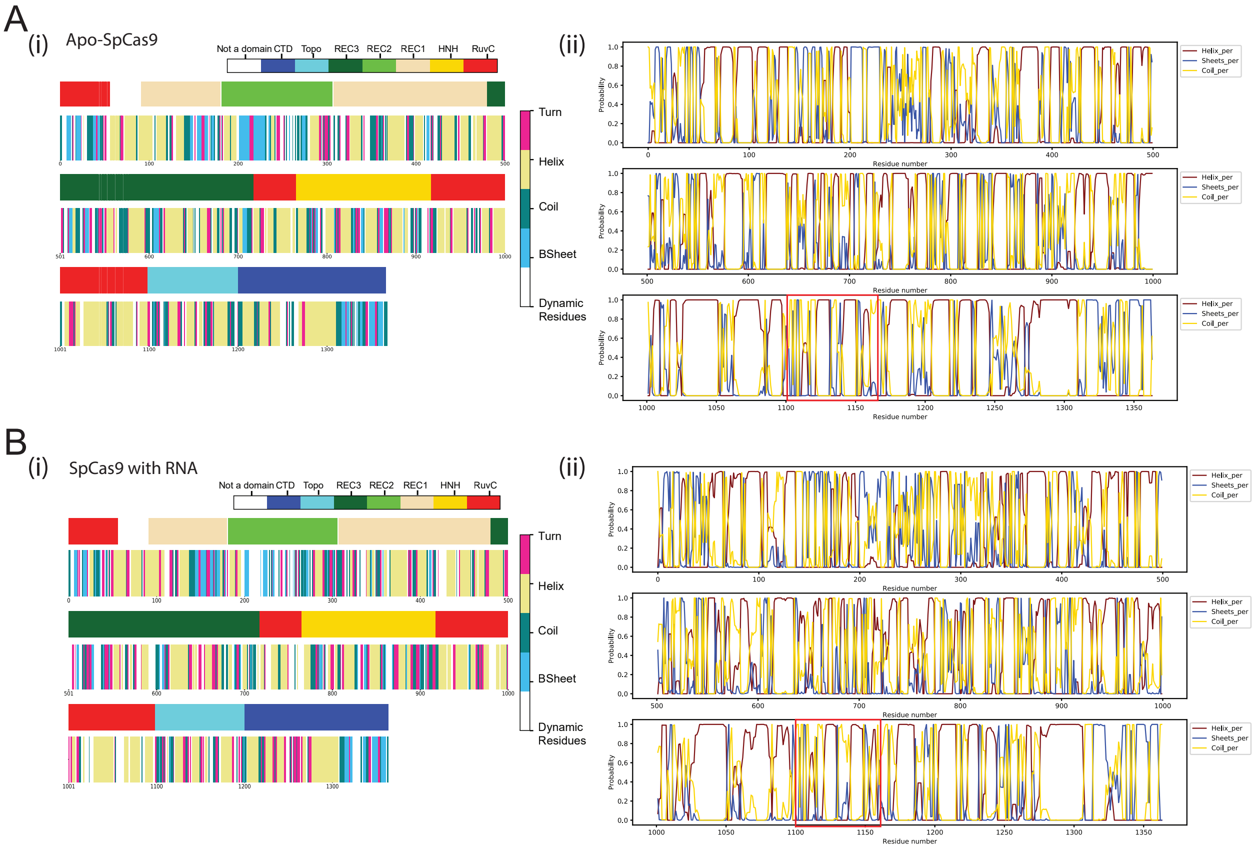


Figure S5: A (i) Residue-wise and domain-wise assignment of secondary structure assignment (present in more than 70% of frames in stable part of trajectory) in Apo-SpCas9. Color coding for domains, RuvC: Red, HNH: Yellow, REC1: Wheat, REC2: Limon, REC3: Dark Green, Topo: Cyan, CTD: Blue and white color for the non-domain region. Color coding for SSE, Helix: Khaki, BSheet: Cyan, Turn: Pink, Coil: Teal and Dynamic Residue: White. (ii) Secondary Structure propensity of every residue in Apo-SpCas9 in 1000 frames. Helix Probability (brown), Sheets probability (blue) and Coil probability (yellow). Regions of higher atomic fluctuations (in RMSF plot) are shown in Red boxes. B (i) Residue-wise and domain-wise assignment of secondary structure assignment (present in more than 70% of frames in stable part of trajectory) in SpCas9 with gRNA (ii) Secondary Structure propensity of every residue in SpCas9-gRNA in 1000 frames. Helix Probability (brown), Sheets probability (blue) and Coil probability (yellow). Regions of higher atomic fluctuations (in RMSF plot) are shown in Red boxes.

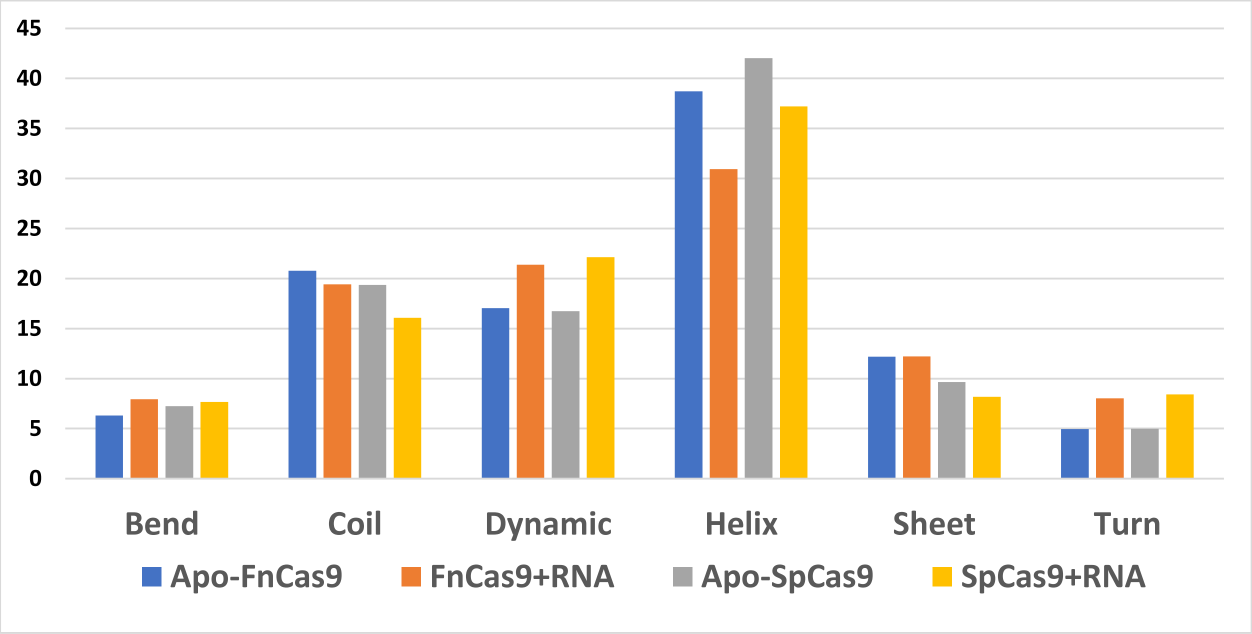


Figure S6: Bar plot showing percentage of Helix, Sheet, Bend, Coil, Turn and dynamic residues in Apo-SpCas9, Spcas9 with gRNA, Apo-FnCas9 and FnCas9 with gRNA.

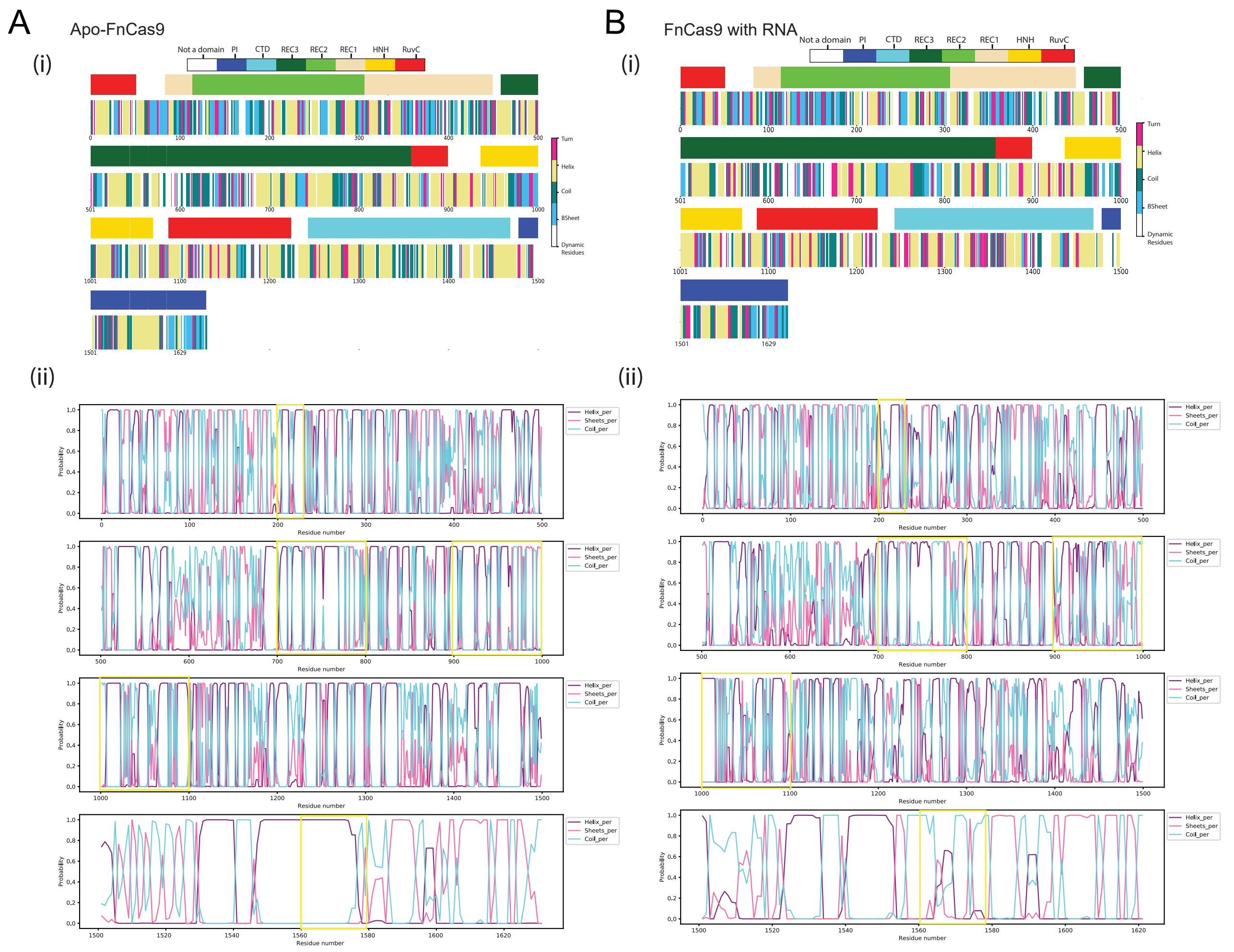


Figure S7: A (i) Residue-wise and domain-wise assignment of secondary structure assignment (present in more than 70% of frames in stable part of trajectory) in Apo-FnCas9. Color coding for domains, RuvC: Red, HNH: Yellow, REC1: Wheat, REC2: Limon, REC3: Dark Green, WED: Cyan, PI: Blue and white color for the non-domain region. Color coding for SSE, Helix: Khaki, BSheet: Cyan, Turn: Pink, Coil: Teal and Dynamic Residue: White. (ii) Secondary Structure propensity of every residue in Apo-FnCas9 in 1000 frames. Helix Probability (violet), Sheets probability (pink) and Coil probability (cyan). Regions of higher atomic fluctuations (in RMSF plot) are shown in yellow boxes. B (i) Residue-wise and domain-wise assignment of secondary structure assignment (present in more than 70% of frames in stable part of trajectory) in FnCas9 with gRNA (ii) Secondary Structure propensity of every residue in FnCas9-gRNA in 1000 frames. Helix Probability (Violet), Sheets probability (pink) and Coil probability (cyan). Regions of higher atomic fluctuations (in RMSF plot) are shown in yellow boxes.

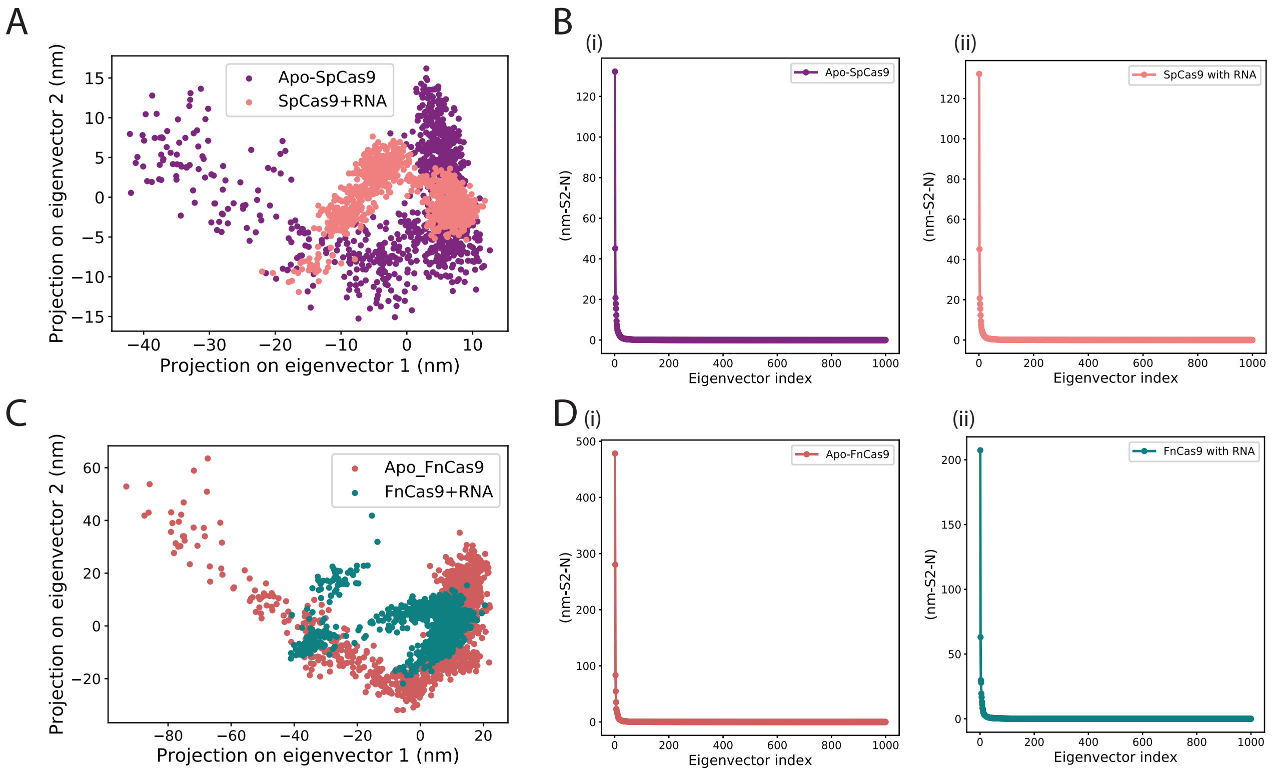


Figure S8: A. 2D projection of backbone atoms in essential subspace along firts two eigenvectors of Apo-SpCas9 (violet) and SpCas9 with gRNA (lightcoral). B. (i) Eigenvalues of the covariance matrix for Apo-Spcas9 (violet). (ii) Eigenvalues of the covariance matrix for SpCas9 with gRNA (lightcoral). C. 2D projection of backbone atoms in essential subspace along first two eigenvectors of Apo-FnCas9 (indian-red) and FnCas9 with gRNA (teal). B. (i) Eigenvalues of the covariance matrix for Apo-Fncas9 (violet). (ii) Eigenvalues of the covariance matrix for FnCas9 with gRNA (teal).

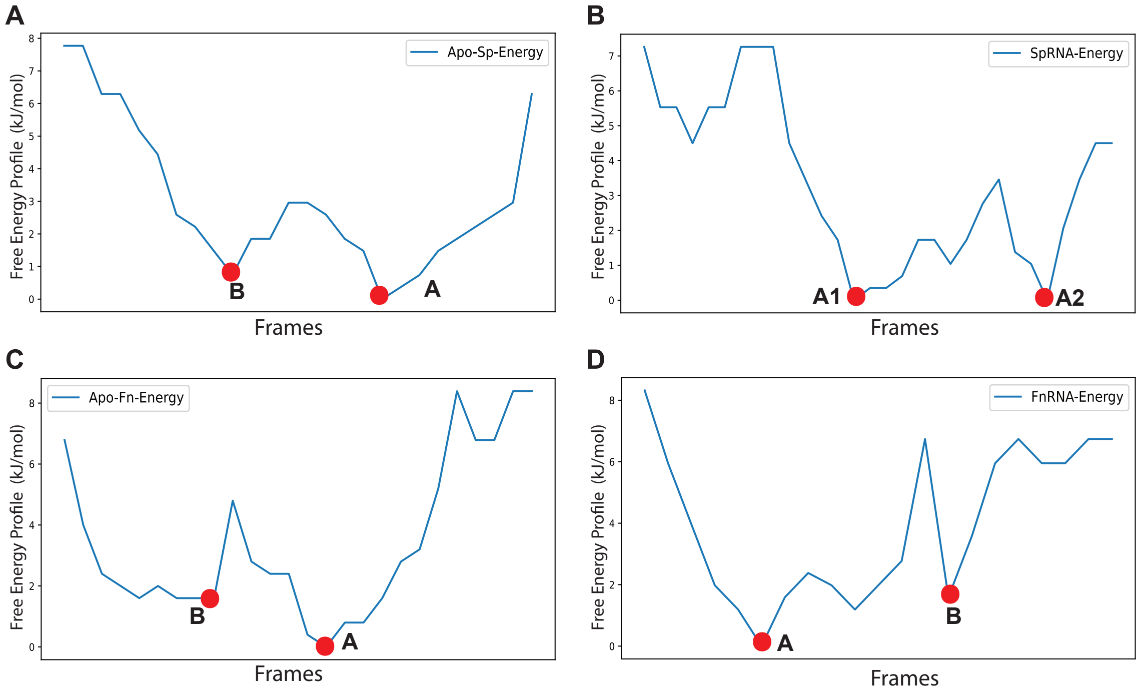


Figure S9: A. Frame-wise Free Energy profile and local energy minimas of A. Apo-SpCas9 B. SpCas9-gRNA bound form. C. Apo-FnCas9 form D. FnCas9-gRNA bound form

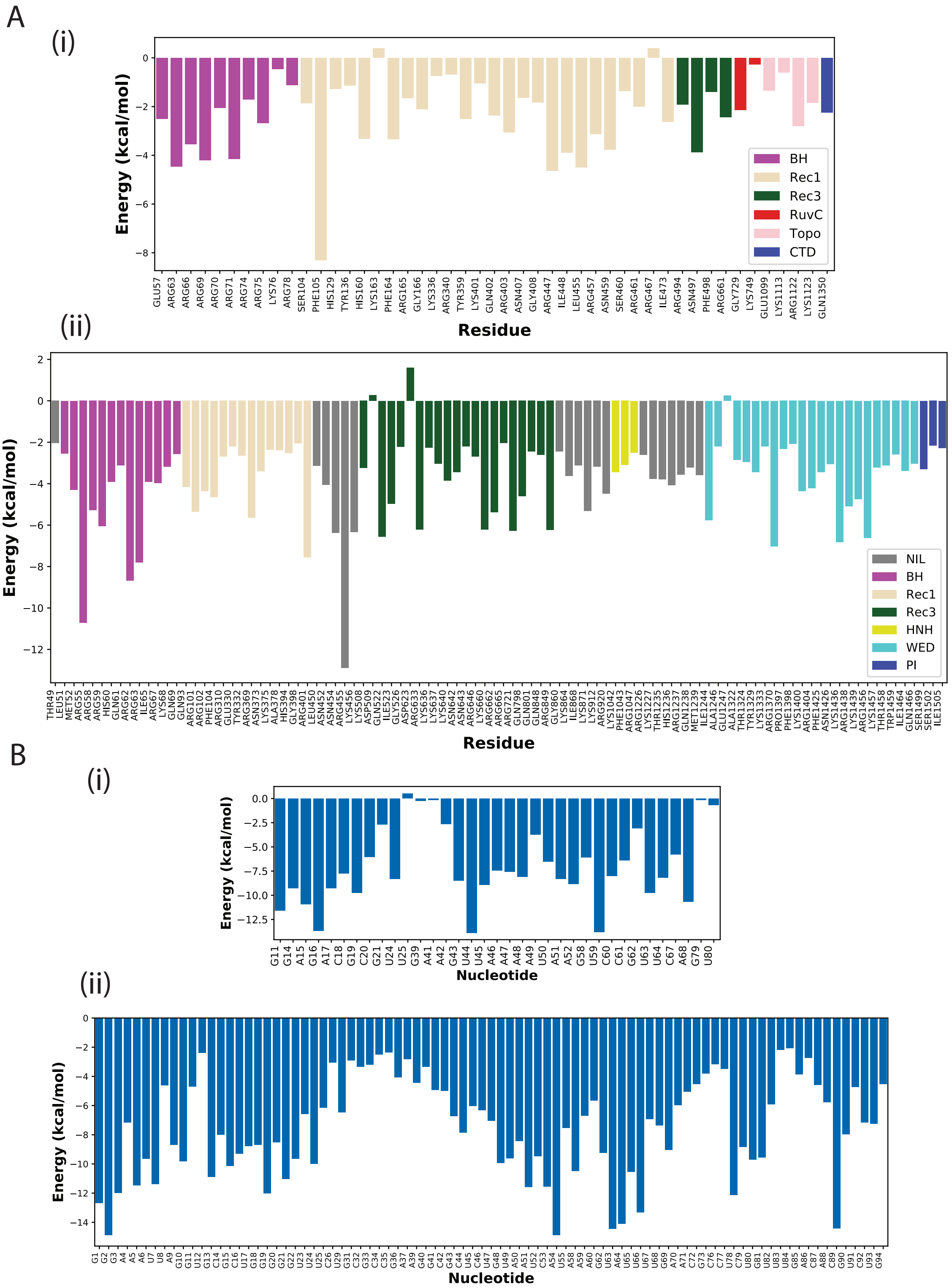


Figure S10: A. Per-residue energy decomposition of residues in SpCas9 protein. (ii) Per-residue energy decomposition of residues in FnCas9 protein. B (i) Per-residue energy decomposition of nucleotides in gRNA bound SpCas9 protein (Energy < -2 kcal/mol). (ii) Per-residue energy decomposition of nucleotides (Energy < -2 kcal/mol) in gRNA bound FnCas9 protein.
