## Supplementary File 2 for "Comparative Structural and Dynamics Study of Free and gRNA-bound FnCas9 and SpCas9 Proteins"

```
CAS9_STRP1  ............................................................
CAS9_FRATN  MNFKILPIAIDLGVKNTGVFSAFYQKGTSLERLDNKNGKVYELSKDSYTLLMNNRTARRH

                                      1        10        20
CAS9_STRP1  ..........................MDKKYSIGLDIGTNSVGWAVITDEYK......VP
CAS9_FRATN  QRRGIDRKQLVKRLFKLIWTEQLNLEWDKDTQQAISFLFNRRGFSFITDGYSPEYLNIVP

            30         40        50        60        70         80
CAS9_STRP1  SKKFKVLGNT.DRHSIKKNLIGALLFDSGETAEATRLKRTARRRYTRRK.NRIC......
CAS9_FRATN  EQVKAILMDIFDDYNGEDDLDSYLKLATEQESKISEIYNKLMQKILEFKLMKLCTDIKDD

                         90       100           110       120
CAS9_STRP1  .....YLQEIFSNEMAKVDDSFFHRLEESFLVEE....DKKHERHPIFGNIVDEVAY..H
CAS9_FRATN  KVSTKTLKEITSYEFELLAD.YLANYSESLKTQKFSYTDKQ.......GNL.KELSYYHH

          130       140             150       160       170
CAS9_STRP1  EKYPTIYHLRK......KLVDS..TDKADLRLIYLALAHMIKFRGHFLIEGDLNPD..NS
CAS9_FRATN  DKYNIQEFLKRHATINDRILDTLLTDDLDIWNF.....NFEKF......DFDKNEEKLQN

          180       190       200                   210       220
CAS9_STRP1  DVDKLFIQLVQTYNQLFEENPIN...ASG........VDAKAILSARLSKSRRLENLIAQ
CAS9_FRATN  QEDKDHIQ.AHLHHFVFAVNKIKSEMASGGRHRSQYFQEITNVLDENNHQEGYLKNFCEN

           230               240       250       260            270
CAS9_STRP1  LPGEK......KN..GLFGNLIALSLGLTPNFKSNFDLAE.....DAKLQLSKDTYDDD.
CAS9_FRATN  LHNKKYSNLSVKNLVNLIGNL............SNLELKPLRKYFNDKIHAKADHWDEQK

                        280                 290              300
CAS9_STRP1  .........IDNLLAQIGDQ.........YADLFLAAK......NLSDAILLSDILR...
CAS9_FRATN  FTETYCHWILGEWRVGVKDQDKKDGAKYSYKDLCNELKQKVTKAGLVDFLLELDPCRTIP

              310       320                                   330
CAS9_STRP1  ..VNTEITKAPLSASMI.............KRY..............DEHHQDLTLLKAL
CAS9_FRATN  PYLDNNNRKPPKCQSLILNPKFLDNQYPNWQQYLQELKKLQSIQNYLDSFETDLKVLKSS

           340                                350       360       370
CAS9_STRP1  VRQQLPEKYK.....................EIFFDQSKNGYAGYIDGGASQE....EFY
CAS9_FRATN  KDQPYFVEYKSSNQQIASGQRDYKDLDARILQFIFDRVK..........ASDELLLNEIY

                         380         390       400         410       420
CAS9_STRP1  KFIKPI.......LEKMDGT...EELLVKLNREDLLRKQRT...FDNGSIPHQIHLGELH
CAS9_FRATN  FQAKKLKQKASSELEKLESSKKLDEVIANSQLSQILKSQHTNGIFEQGTF........LH

                    430       440       450                    460
CAS9_STRP1  AILRRQEDFYPFLKDNREKIEKILTFRIPYY...........VGPLARGNSRFAWMTRKS
CAS9_FRATN  LVCK....YY....KQRQRARDSRLYIMPEYRYDKKLHKYNNTGRFDDDNQLLTYCNHKP
```

```
                 470                                 480              490                  500            510
                   .                                   .                .                    .              .
CAS9_STRP1  EE.............TITPWNFEEVVDK.GASAQSFI.....ERMTNFDKNLPNEKVLPK
CAS9_FRATN  RQKRYQLLNDLAGVLQVSP.NF..LKDKIGSDDDLFISKWLVEHIRGFKKACEDSLKIQK

                                                   520              530              540
                                                     .                .                .
CAS9_STRP1  .......HSLLYEYFT.......VYNELTKVKYVTEGMRKPAFLSGEQKKA.........
CAS9_FRATN  DNRGLLNHKINIARNTKGKCEKEIFNLICKI....EG........SEDKKGNYKHGLAYE

             550              560              570                         580              590              600
               .                .                .                           .                .                .
CAS9_STRP1  IVDLLFKTNRKVTVKQLKEDYFKKIECFDSV.......EISGVEDRFNASLGTYHDLLKI
CAS9_FRATN  LGVLLFGEPNEAS....KPEFDRKIKKFNSIYSFAQIQQIAFAERKGNANTCA.......

                          610              620              630              640              650              660
                            .                .                .                .                .                .
CAS9_STRP1  IKDKDFLDNEENEDILEDIVLTLTLFEDREMIEERLKTYAHLFDDKVMKQLKRRRYTGWG
CAS9_FRATN  ......VCSADNAHRMQQIKIT.......EPVEDN........KDKIILSAKAQRLPA..

                          670                   680              690              700              710
                            .                     .                .                .                .
CAS9_STRP1  RLSRKLINGIRDKQS...GKTILDFLKSDGFANRNFMQLIHDDSLTFKEDIQKAQVSGQG
CAS9_FRATN  .IPTRIVDGAVKKMATILAKNIVD....DNW..QNIKQVL....................

             720                 730              740                   750              760              770
               .                   .                .                     .                .                .
CAS9_STRP1  DSLHE.HIANLAGSPAIK.KGILQTVK...VVDELVKVMGRHKPENIVIEMARENQTTQK
CAS9_FRATN  SAKHQLHIPIITESNAFEFEPALADVKGKSLKDRRKKALERISPENIF............

                       780              790              800              810              820              830
                         .                .                .                .                .                .
CAS9_STRP1  GQKNSRERMKRIEEGIKELGSQILKEHPVENTQLQNEKLYLYYLQNGRDMYVDQELDINR
CAS9_FRATN  ..KDKNNRIKEFAKGISAYSGANLTD..................................

                              840              850                   860                             870
                                .                .                     .                               .
CAS9_STRP1  LSDYD.....VDHIVPQSFLKDDSIDNKV....LTRSD.KNRGK........SDNVPSE.
CAS9_FRATN  .GDFDGAKEELDHIIPRSHKKYGTLNDEANLICVTRGDNKNKGNRIFCLRDLADNYKLKQ

                                 880              890              900              910              920
                                   .                .                .                .                .
CAS9_STRP1  .......EVVKKMKN.YW.RQLLNAKLITQRKFDNLTKAERGGLSELDKAGFIKRQLVET
CAS9_FRATN  FETTDDLEIEKKIADTIWDANKKDFKFGNYRSFINLTPQEQ.......KAFRHALFLADE

                    930              940              950                 960                   970
                      .                .                .                   .                     .
CAS9_STRP1  RQITKHVAQILDSRMNTKYDENDKLIREVKV..ITLKSK...LVSDFRKDFQFYKVREIN
CAS9_FRATN  NPIKQAVIRAINNRNRTFVNGTQRYFAEVLANNIYLRAKKENLNTD.KISFDYFGIPTIG

               980              990             1000                1010             1020             1030
                 .                .                .                   .                .                .
CAS9_STRP1  NYHHAHDAYLNAVVGTALIKK.YPKLESEFVYGDYKVYDVRKMIAKSEQEIGKATAKYFF
CAS9_FRATN  NGR...........GIAEIRQLYEKVDS.....DIQAY......AKGD....KPQASYSH

              1040             1050             1060             1070               1080             1090
                 .                .                .                .                  .                .
CAS9_STRP1  YSNIMNFFKTEITLANGEIRKRPLIETNGETG.EIVWDKGRDFATVRKVLSMPQVNIVKK
CAS9_FRATN  LIDAMLAF....CIADEHR......NDGSIGLEI..DKNYSLYPLDK..........NT
```

```
           1100      1110      1120      1130                      1140
CAS9_STRP1  TEVQTGG.FSKESILPKRNSDKLIARKKDWDPKKYGGFDS...............PTVAY
CAS9_FRATN  GEVFTKDIFSQIKITDNEFSDKKLVRKKAIE.....GFNTHRQMTRDGIYAENYLPILIH

                 1150       1160      1170      1180        1190
CAS9_STRP1  SVLVVAKVEKGKS.KKLKSVKELLGITIMERSSFEKNPIDFLEAKGY..KEVKKDLIIKL
CAS9_FRATN  KEL..NEVRKGYTWKNSEEIKIFKG.........KKYDIQQLNNLVYCLKFVDKPISIDI

          1200      1210      1220      1230      1240      1250
CAS9_STRP1  PKYSLFELENGRKRMLASAGELQKGNELALPSKYVNFLYLASHYEKLKGSPEDNEQK..Q
CAS9_FRATN  QISTLEELRN...........ILTTNNIAATAEY........YYINLK......TQKLHE

            1260                          1270      1280        1290
CAS9_STRP1  LFVEQ......HKHYLDEI..............IEQISEFSKRVILADAN..LDKVLSAY
CAS9_FRATN  YYIENYNTALGYKKYSKEMEFLRSLAYRSERVKIKSIDDV.KQVLDKDSNFIIGKITLPF

              1300      1310      1320      1330        1340      1350
CAS9_STRP1  NKHRDKPIREQAENIIHLFTLTNLGAPAAFKYFDTTI..DRKRYTSTKEVLDATLIHQSI
CAS9_FRATN  KKEWQRLYRE.....................WQNTTIKDDYEFLKSFFNVKSITKLHKKV

                1360
CAS9_STRP1  TGLYETRIDLSQLGGD............................................
CAS9_FRATN  RKDFSLPISTNE..GKFLVKRKTWDNNFIYQILNDSDSRADGTKPFIPAFDISKNEIVEA

CAS9_STRP1  ............................................................
CAS9_FRATN  IIDSFTSKNIFWLPKNIELQKVDNKNIFAIDTSKWFEVETPSDLRDIGIATIQYKIDNNS

CAS9_STRP1  ............................................................
CAS9_FRATN  RPKVRVKLDYVIDDDSKINYFMNHSLLKSRYPDKVLEILKQSTIIEFESSGFNKTIKEML

CAS9_STRP1  ..............
CAS9_FRATN  GMKLAGIYNETSNN
```
